# Constructing a Mouse Brain Atlas of Dendritic Microenvironments Helps Discover Hidden Associations Between Anatomical Layout, Projection Targets and Transcriptomic Profiles of Neurons

**DOI:** 10.1101/2024.09.22.614330

**Authors:** Yufeng Liu, Sujun Zhao, Zhixi Yun, Feng Xiong, Hanchuan Peng

## Abstract

Digital brain atlases have become essential anatomical references for understanding the spatial and functional organization of brains. For mice, typical resources include the Allen Reference Atlas, the Allen Common Coordinate Framework (CCF), and their variants, like CCFv3. However, previous whole-brain atlases were constructed based on limited neuronal features, such as cell body (soma) density or average maps from collections of registered brain images, without considering the spatial organization of neuronal arbors. This study introduces a microenvironment representation that incorporates the morphological features of neighboring neurons to better quantify brain modularity. We generated a large dataset containing dendrites from 101,136 neurons across 111 mouse brains, covering 91% of non-ventricular, non-fiber-tract CCF regions, and constructed a multidimensional microenvironment feature map of the whole brain. Our findings reveal that the spatial organization of these microenvironments outperforms the CCFv3 and a state-of-the-art spatial transcriptomic cell atlas by providing complementary subregions within established regions, nearly doubling the total number of brain regions compared to CCFv3. In this way, our atlas enables the identification of previously unobserved neuron groupings or “subtypes”. Our results also demonstrate that this microenvironment atlas enhances local spatial homogeneity while maintaining spatial differentiation within established CCF brain regions. For example, we found that the microenvironments of hippocampal neurons are correlated with axonal projection targets and improve the specificity of projection mapping, which implies the potential characterization of long-range axonal projections of mammalian neurons based on only local dendritic organization. The sub-parcellation of the caudoputamen (CP) aligns well with previous studies on projections, connectivity, and transcriptomics, revealing diverse input and output wiring patterns among CP subregions.

## Introduction

Digital brain atlases are critical resources by providing a consistent anatomical framework for cross-brain mapping that facilitates the integration of diverse datasets and enables researchers to compare findings across studies. For example, the Allen Brain Atlas (Dong, 2008; Hawrylycz et al., 2012) has been pivotal in advancing the understanding of gene expression patterns and their relationships to brain anatomy and function. The Common Coordinate Framework (CCF) version 3 (Wang et al., 2020) serves as a valuable reference for aligning neuroimaging and neuroanatomical data. These atlases also spur the development of new hypotheses and the discovery of previously unobserved cell types and subtypes (Zeisel et al., 2015; Codeluppi et al., 2018).

Most existing mouse brain atlases are constructed upon collections of images capturing neuronal features like histological stains, gene expression and connectivity, as seen in the Allen Reference Atlas (Dong, 2008) and the Franklin-Paxinos atlas (Franklin & Paxinos, 2008), or combinations of multimodal features in CCFv3 (Wang et al., 2020). One of the major limitations of these approaches is the inadequate resolution in certain brain regions, making it difficult to capture fine-grained anatomical details. This challenge arises from the limited discriminative power of the utilized neuroanatomical features, such as cell density. Indeed, there has been debate among neuroanatomists on the accuracy and granularity of several CCF regions (Chon et al., 2019), which call for independent validation using alternative labeling and annotation standards. This problem is even more obvious when various levels of anatomical deformation are introduced in different sample preparation methods used in developing these atlases, such as serial thick sections (e.g., serial two-photon tomography in TissueCyte; Ragan et al., 2012), serial thin sections (e.g., fluorescence micro-optical sectioning tomography (fMOST); Gong et al., 2013, 2016), and other tissue-clearing techniques (e.g., CLARITY; Chung et al., 2013) and tissue-expansion methods (e.g., expansion microscopy; Murakami et al., 2018).

A valuable approach for constructing a high-resolution brain atlas is to incorporate more functionally informative features, such as single-cell transcriptomics. Single-cell spatial transcriptomic techniques like 10x Genomics and multiplexed error-robust fluorescence *in situ* hybridization (MERFISH; Chen et al., 2015) have recently gained popularity in creating systematic cell atlases (Yao et al., 2023; Zhang et al., 2023). However, studies on the mouse brain have shown that the cellular organization revealed by MERFISH does not always align with the anatomical layout of the CCFv3 atlas (Yao et al., 2023). Therefore, building a high-resolution brain atlas remains a challenge.

Here, we provide an orthogonal approach to the above efforts. We notice that single-neuron morphology has long been recognized as a critical feature for determining neuron types and brain anatomy (Zeng & Sanes, 2017; Zeng, 2022). Although obtaining large-scale full morphologies remains difficult, dendritic morphologies are more readily available. However, dendritic morphologies are often stereotyped across populations, sparking debates about their ability to differentiate neuron types (Jiang et al., 2015). In our previous research, we quantified the stereotypy and diversity of dendritic features across cortical regions in the human brain (Han et al., 2023). We extended this approach to develop whole-brain dendritic microenvironments using the 3D dendritic morphologies of over 100,000 neurons in mouse brains. By integrating the morphological features of neighboring neurons, these microenvironments enhance neuroanatomical discrimination and show correlations with axonal projection patterns. This framework also facilitates the creation of a high-resolution mouse brain atlas, CCF-ME, enabling the detailed characterization of brain anatomy, projection specificity, and transcriptional patterns with greater granularity.

## Results

### A high-resolution dendritic microenvironment atlas of the mouse brain

We developed a high-resolution whole-brain atlas for the mouse by analyzing dendritic microenvironments, achieving greater granularity than existing brain atlases. This new atlas is based on the auto-reconstruction of 101,136 neurons (**Figure 1A**) from 111 sparsely labeled fMOST mouse brains (Peng et al., 2023; **Supplementary Table S1**). Each neuron was reconstructed from a ∼256 µm width image volume centered on the cell body, capturing the majority of the dendrites. These neurons span all seven major brain regions: cerebral cortex (CTX), cerebral nuclei (CNU), midbrain (MB), thalamus (TH), cerebellum (CB), hindbrain (HB), and hypothalamus (HY), with particularly high representation in CTX (46.2%) and CNU (30.1%) (**Supplementary Figure S1B-C**). These neurons span the majority of brain regions (528 out of 582 non-fiber tract, non-ventricle salient regions in the CCFv3 atlas), with an average of 192 neurons per region. Among them, 397 regions host at least ten neurons each.

**Figure 1.**
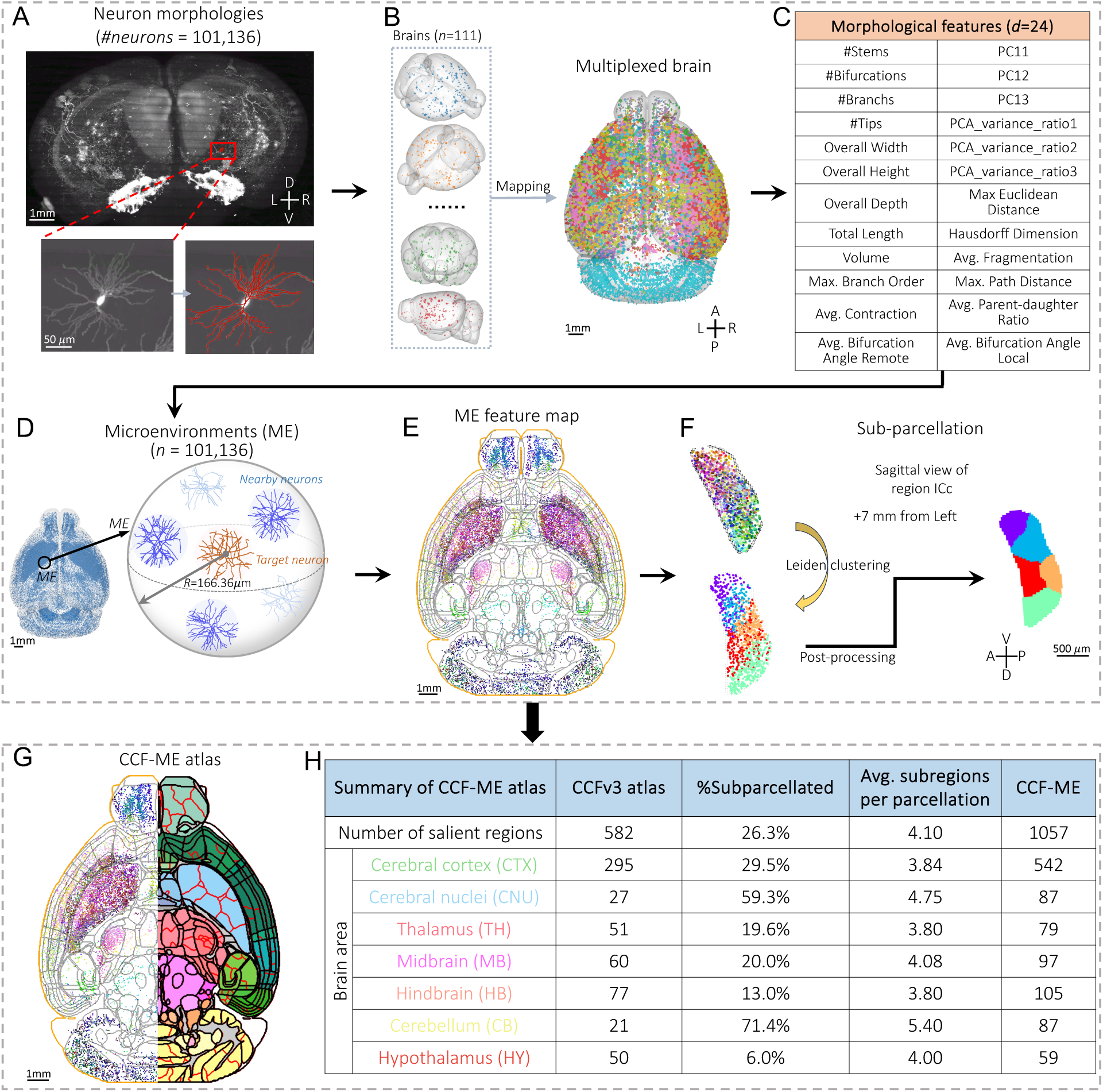
Overview of sub-parcellation and the CCF-ME atlas. **A**. Coronal view of a sparsely labeled mouse brain with an exemplar neuron highlighted. The auto-traced morphology of the neuron is overlaid on the image (bottom right). D, V, L, and R: dorsal, ventral, left, and right. **B**. A multiplexed brain containing all morphologies mapped from the original brains. Morphologies are colored according to the brains they come from. A, P, L, and R: anterior, posterior, left, and right. **C**. The morphological features utilized in this work (see **Methods**). **D**. Construction of a microenvironment. **E**. The horizontal middle slice with microenvironments within 0.5 mm projected onto the slice. Microenvironments are RGB-colored by the top three morphological features identified using the mRMR algorithm. The region boundaries are outlined with gray dots, and the outermost boundary of the brain is highlighted in orange. **F**. An example of sub-parcellation of the region ICc based on the microenvironment features. Different colors on the right mask indicate different subregions. A, P, D, and V: anterior, posterior, dorsal, and ventral. **G**. Illustration of the CCF-ME atlas. Left, the left half of the horizontal middle slice of the microenvironment feature map. Right, CCF-ME, with new boundaries highlighted in red line. **H**. A summary table comparing CCF-ME and CCFv3. The column “Avg. subregions per parcellation” shows the average number of subregions in CCF-ME identified for each subdivided CCF region. Only sub-parcellated CCF regions are considered when calculating “Avg. subregions per parcellation”.

Each neuron was mapped to the CCFv3 atlas, placing them in a standardized isotropic coordinate space (**Figure 1B**). For comparability, we extracted the soma-connected subtree within a 100 µm radius around the cell body for each neuron and represented it by a 24-dimensional morphological feature vector (**Figure 1C**; **Methods**). The auto-traced neurons (examples in **Supplementary Figure S1D**) exhibit a median bi-directional distance of 1.926 µm compared to manual annotations (**Supplementary Figure S1F**), with morphological features similar between the two methods, aside from minor variances (**Supplementary Figure S1E**). Due to inherent differences in data generation and skeleton sparsity, local features may diverge between manual reconstructions and auto-traced morphologies. These biases are evident in the slight decreases observed in local bifurcation angle (“Avg. Bifurcation Angle Local”) and branch length (“Avg. Fragmentation”; **Supplementary Figure S1E**). The small variances of these features indicate that they should be biased rather than erroneous (**Supplementary Figure S1E**).

We constructed microenvironments for each neuron by summarizing the morphological features of up to six neighboring neurons (**Figure 1D**; **Supplementary Figure S2A**). This representation is inspired by previous morpho-anatomical analyses of the human brain (Han et al., 2023). The microenvironments not only helped mitigate possible tracing flaws but also enhanced discriminative power between different neuron types, as demonstrated by higher silhouette scores for K-Means clustering compared to single neuron dendrites (**Supplementary Figure S2B**). Using the Leiden community detection algorithm, we classified microenvironments into communities based on their nearest neighbors, followed by post-processing to generate smooth subregion boundaries (**Supplementary Figure S2C**).

To explore morphological distributions across the brain, we created a whole-brain feature map using three key features: total neurite length (“Total Length”), average straightness of branch (“Avg. Contraction”), and branch length (“Avg. Fragmentation”, defined as the number of 2 µm-long compartments in a branch). These features were selected using the minimal-Redundancy-Maximal-Relevance (mRMR; Peng et al., 2005) algorithm. We color-coded microenvironments based on these three features, assigning RGB values to each feature respectively. The resulting map highlights spatial coherence and reveals inter- and intra-regional variations (**Figure 1E**), such as the spatial differentiation observed in neurons of the inferior colliculus (ICc): ventral neurons exhibit longer total length (red points; **Figure 1F**), dorsal neurons show more straight branches (green points; **Figure 1F**). As a result, ICc neurons are segmented into five subregions (**Figure 1F**).

From these dendritic microenvironments, we constructed a high-resolution atlas, CCF-ME, comprising 1,057 non-ventricular, non-fiber-tract salient regions, nearly double the 582 regions in CCFv3 (**Figure 1G**). In CCF-ME, cortical regions remain consistent (51.3% vs. 50.8%), while subregions in cerebral nuclei and the cerebellum increased significantly by 78% and 128%, respectively (**Supplementary Figure S3A)**, indicating a higher number of parcellated subregions in these brain areas (**Supplementary Figure S3B**). Approximately 26.3% (*n*=153) of CCF regions were subdivided into an average of 4.1 subregions (**Figure 1H**). The regions in CB exhibit the highest average number of subregions among common brain areas and a high ratio of region sub-parcellation (5.4 and 71.4%; **Figure 1G-H**), whereas only 6% of HY regions are sub-parcellated (**Figure 1H**). Thalamus (TH) and hindbrain (HB) regions show a slightly higher ratio of sub-parcellated CCF regions (19.6 and 13%; **Figure 1H; Supplementary Figure S3B**).

We examined four metrics: number of neurons, volume, feature variability (“Feature STD”), and spatial autocorrelation (“Moran’s Index”), and found that volume and neuron count positively correlated with the number of subregions (**Supplementary Figure S3C**, left). Non-linear relationships were observed for feature variance and Moran’s Index, where regions with intermediate values of these metrics exhibited the highest subregion counts (**Supplementary Figure S3C**, right). This is reasonable as a small Moran’s Index and feature variance value indicates a highly intermingled morphology distribution, and a large positive Moran’s Index value represents a homogeneous distribution, both of which preclude the subdivision of a region. As a whole, the number of parcellated subregions correlates to all metrics considered, but in different manners.

As expected, the regions in the CCF-ME atlas have smaller volumes (**Supplementary Figure S3D**). The average volume in CCF-ME is 0.21 mm³, smaller than that in the CCF atlas (0.39 mm³). Only 3% of regions in CCF-ME have a volume larger than one mm³, compared to 8.6% in the CCFv3 atlas. The proportion of regions with a small volume (> 0.027 mm³ but < 0.2 mm³) increased from 40.9% in CCFv3 to 59.7% in CCF-ME (**Supplementary Figure S3D**). For each CCF region that was divided into subregions, their volumes showed a modest uniformity, with a Gaussian distribution (*μ*=0.28) of the Gini coefficients of subregion volumes (**Supplementary Figure S3F**).

### Microenvironments improve local spatial coherence and reveal subregional differentiation

The large-scale dendritic reconstructions allow for detailed characterization of morphology and anatomical organization. By employing an ensemble-based microenvironment representation, the local coherence of morphological features was enhanced, revealing spatial differentiation throughout the brain. To illustrate, we compared the spatial distribution of microenvironments to soma distribution and single-neuron dendrite distribution. We analyzed seven coronal slices ranging from 1.5 mm to 11.5 mm along the anterior-posterior (AP) axis (A1-A7; **Figure 2B**). For each slice, neurons within 0.5 mm on either side of the AP axis were color-coded by the top three morphological features and projected onto the slice (**Figure 2A**, left). We also mapped the soma distribution onto the same slices (**Figure 2A**, right) for comparison. These cells were collected from 111 brains labeled by 30 reporter genes (**Supplementary Table S1**).

**Figure 2.**
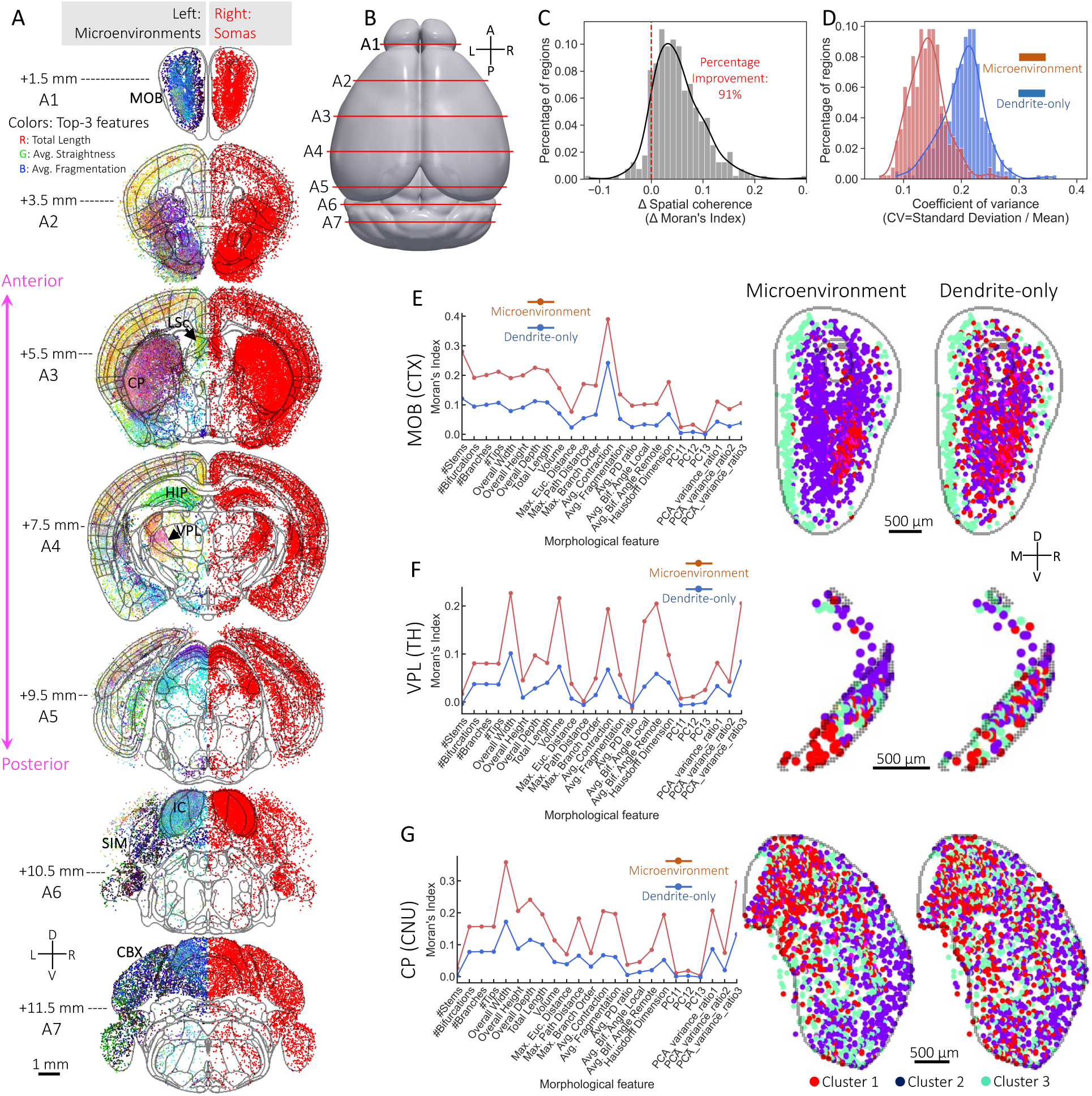
Microenvironment feature maps. **A**. Coronal slices of microenvironments (left) and somas (right) ranging from 1.5 mm to 11.5 mm along the AP axis. A microenvironment is colored based on the values of the top three features, with the feature values assigned to the red, green, and blue channels of the corresponding voxel. To facilitate visualization, histogram equalization is applied to each channel. Somas and microenvironments within 0.5 mm of each slice are projected onto it using maximum intensity projection. The boundaries of each slice are highlighted by gray curves. Brain regions cited in the main text are annotated. D, V, L, and R: dorsal, ventral, left, and right. **B**. Illustration of the location of the seven slices on the CCFv3 atlas. A, P, L, and R: anterior, posterior, left, and right. **C**. The distribution of spatial coherence (calibrated by Moran’s Index score) changes for CCF regions after using microenvironment representation. The vertical red dashed line highlights the location of non-improvement. **D**. The coefficients of variance distributions of single neuron dendrites and microenvironments. **E-G.** Spatial coherence of morphological features (Moran’s Index, first column), coronal slices of microenvironment feature maps (second column), and coronal slices of single neuron dendrite feature maps (third column) for MOB (**E**), VPL (**F**), and CP (**G**). The feature maps are colored based on their clusters estimated using spectral clustering (*k*=3) on the 2D Uniform Manifold Approximation and Projection (UMAP) feature spaces.

Microenvironments showed clear spatial preferences: neurons in the main olfactory bulb (MOB) had smaller total skeleton lengths and more curved branches (**Figure 2A1**), while cortical neurons generally exhibited longer dendrites with straighter, shorter branches (**Figures 2A2-4**). In contrast, hippocampal (HIP) neurons displayed straighter branches (**Figure 2A4**), and distinct patterns emerged in regions like the caudoputamen (CP), inferior colliculus (IC), and cerebellar cortex (CBX) (**Figures 2A3, 2A6, 2A7**). For instance, IC neurons generally have straighter and longer branches, whereas those in surrounding regions, such as the simple lobular (SIM), have more curved branches (**Supplementary Figure S4C**). Systematically, microenvironments generally showed similar dendritic microenvironments within the same brain area, such as the thalamus, where neurons had larger total local dendritic lengths and higher intra-area consistency (**Supplementary Figure S4A-B**). Isocortical neurons display a dispersed total length distribution with modest branch length and straightness (**Supplementary Figure S4A**), resulting in a smaller intra-area consistency (**Supplementary Figure S4B**). Neurons of the cerebellar cortex and cerebellar nuclei have more curved branches and smaller total lengths (**Supplementary Figure S4A**), contributing to considerable intra-area and pairwise consistencies (**Supplementary Figure S4B**). However, neurons within the same area can still display diverse morphologies, as seen in CP and the caudal part of the lateral septal nucleus (LSc) from the striatum (**Figure 2A3**).

Compared to single-neuron dendrites, microenvironments showed improved spatial coherence. Among 397 CCF regions with at least ten neurons, 91% exhibited improved Moran’s Index scores with microenvironments (**Figure 2C**), along with reduced intra-region feature variances (**Figure 2D**). This spatial coherence improvement was evident in clusters in regions such as the main olfactory bulb (MOB) in the cortex (**Figure 2E**), the ventral posterolateral nucleus (VPL) in the thalamus (**Figure 2F**), and the CP in the cerebral nuclei (**Figure 2G**), where microenvironments consistently showed higher feature coherence compared to individual neurons.

The microenvironments illustrated clear spatial differentiation within many CCF regions. clear spatial differentiation within many CCF regions. In particular, microenvironments across various subregions of these regions displayed distinct morphological features, as evidenced by their spatial distributions in regions like MOB (**Figure 2A1; Supplementary Figure S4D**), VPL (**Figure 2A4**) and CP (**Figure 2A3**). For clear visualization, we classified the microenvironments, and the resulting clusters exhibited clear spatial preferences across all regions (**Figure 2E-G**). Another example is the nucleus accumbens (ACB), where neuronal morphologies were distributed in a structured manner (**Supplementary Figure S4D**).

In summary, microenvironments enhance local spatial coherence of neuronal morphological layout while preserving spatial differentiation at the subregion level by integrating the morphological features of neighboring neurons. This approach allows the spatial organization of whole-brain morphologies to be inferred from microenvironments and facilitates subregional parcellations within established CCF brain regions.

### Microenvironments correlate with axonal projection of hippocampal neurons

Given the spatial preference and subregional differentiation observed in microenvironments (**Figure 2**), an important question arises: what is the relationship between dendritic morphologies, microenvironments, and axonal projection patterns? Previous research has reported that dendritic and axonal patterns can be consistent in some prefrontal cortical subtypes while divergent in others (Gao et al., 2023). We aimed to explore this relationship using a recently released hippocampus dataset (Qiu et al., 2024), focusing on 3,822 neurons with dendrites reconstructed and at the same time, their somas were located within the eight hippocampal formation regions — CA1, CA2, CA3, subiculum (SUB), prosubiculum (ProS), and the molecular, polymorph, and granule cell layers of the dentate gyrus (DG-mo, DG-po, DG-sg) (**Supplementary Figure S5A**).

The microenvironments of hippocampal neurons revealed three distinct clusters (α, β, and γ; **Figure 3B**), in contrast to the less distinguishable single-neuron dendritic morphologies (**Figure 3A**), demonstrating a more divergent yet internally prototyped pattern when using the microenvironment representation. The spatial layout of these clusters aligns well with the CCFv3 hippocampal anatomy on all six coronal slices (**Figure 3D**). Specifically, cluster γ neurons are primarily located in CA3, cluster α in DG-sg, while neurons in SUB, ProS, and CA1 fall into cluster β (**Figure 3D**; **Figure 3G**).

**Figure 3.**
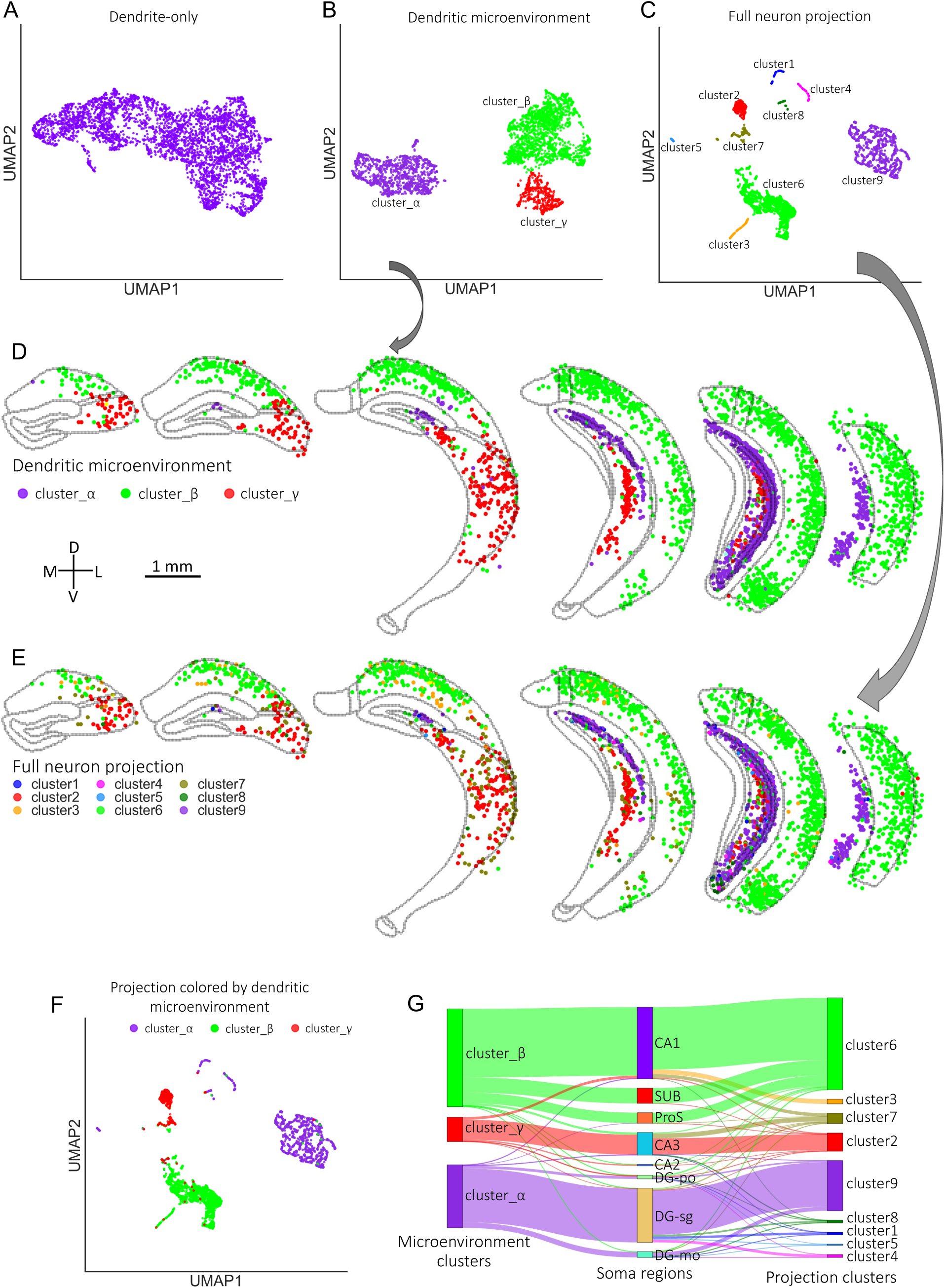
Correspondence between microenvironments and axonal projections of hippocampal neurons. **A-C**. UMAP visualization of clusters for single-neuron dendritic morphologies (**A**), dendritic microenvironments (**B**), and full neuron projections of hippocampal neurons (**C**). The projection of a neuron was a region-wise vector, where each item was the projection intensity in a target region. The projection intensity was estimated as *ln*(length+1), where “length” is the total axonal length in a target region. Regions with total axonal length < 1 mm were discarded. **D-E**. Coronal slices showing the spatial distribution of various clusters of microenvironments (**D**) and axonal projections (**E**). The boundaries of brain regions are highlighted with curves. The masks of the regions are displayed in **Supplementary Figure S5A**. D, V, M, and L: dorsal, ventral, medial, and lateral. **F**. UMAP visualization of axonal projections colored by dendritic microenvironment cluster types. **G**. A Sankey diagram illustrating the correspondence between dendritic microenvironment clusters (left), the soma regions in CCFv3 atlas (middle), and the axonal projection clusters (right).

Axonal projections are more divergent and categorized into nine clusters (clusters 1-9; **Figure 3C**). The spatial distribution of projection clusters mirrored the dendritic microenvironments (**Figure 3D-E**). Notably, projection clusters 6, 9, and 2 correspond to microenvironment clusters β, α, and γ, respectively (**Figure 3F-G**). Projection cluster 7 corresponds to both microenvironment clusters β and γ, without a strong bias towards either (**Figure 3F-G**). This correspondence between dendritic microenvironments and axonal projections indicates that dendritic microenvironments may be capable of predicting long axonal projection patterns, a significant insight given the difficulty of reconstructing long-projecting axons in large quantities.

The finding should also hold for microenvironments constructed upon auto-traced morphologies, which showed similar feature values and low reconstruction errors (**Supplementary Figure S1**). At the same time, the spatial distribution of microenvironment features was consistent with that of manually annotated neurons in colocalized slices (slices B5 and B6; **Supplementary Figure S5B5-6; Supplementary Figure S5C5-6**). For instance, medial neurons were colored green, while lateral neurons appeared in magenta for both their microenvironments and the manually annotated neurons in slice 5 and slice 6 (**Supplementary Figure S5B5-6; Supplementary Figure S5C5-6**). Meanwhile, the multidimensional spatial organization of hippocampal regions along the longitudinal and transverse axes is also reported in previous experiments utilizing genomic, anatomical, and functional techniques (Thompson et al., 2008; Dong et al., 2009; Fanselow & Dong, 2010; Strange et al., 2014; Bienkowski et al., 2018). Stretching the slices along the longitudinal axis further highlighted these layered differentiation and spatial patterns in both datasets (**Supplementary Figures S5D; Supplementary Figures S6**). Even when up to 40 skeletal compartments and subsequent branches (41% of the total skeleton length of a neuron) were removed, the spatial layout of microenvironments remained robust, with only minor decreases in Moran’s Index from 0.51 to 0.44 (**Supplementary Figure S7**). The deterioration is significantly more severe than in our auto-traced local dendrites, which retained a median length of 93% compared to the manually annotated dendrites (dashed red line in **Supplementary Figure S7A; Supplementary Figure S1E**). The resilience of spatial patterns, even under significant perturbation, reinforces the robustness of the microenvironments based on auto-traced morphologies.

### Microenvironments improve axonal projection mapping specificity

The correlation between microenvironments and axonal projection patterns in hippocampal neurons suggests that CCF-ME may enhance projection mapping specificity. To demonstrate, we compared the projection patterns of manually annotated hippocampal neurons (Qiu et al., 2024) across the CCFv3 atlas and our CCF-ME atlas.

Neurons from different CCF-ME subregions within the same CCF region exhibited distinct projection patterns. For example, neurons in subregion R7 of CA3 demonstrated lower overall projection intensity and projected exclusively to ipsilateral targets, including subregions CA1-R1, CA3-R2, CA3-R7, and DG-po-R1 (**Figure 4A**). In contrast, neurons from subregion R4 of the presubiculum (PRE) predominantly projected to contralateral hippocampal subregions (**Figure 4A**). Additionally, hippocampal neurons showed clear local projection preferences, characterized by enriched ipsilateral connections within hippocampal subregions (**Figure 4A**, cyan rectangles). Other subregions, such as those in CA3, also exhibited unique, subregion-specific projection patterns (**Figure 4A**). These findings highlight the increased specificity of projection mapping when using the CCF-ME atlas (**Figure 4A**, right), which was not discernible in the CCFv3 atlas (**Figure 4A**, left).

**Figure 4.**
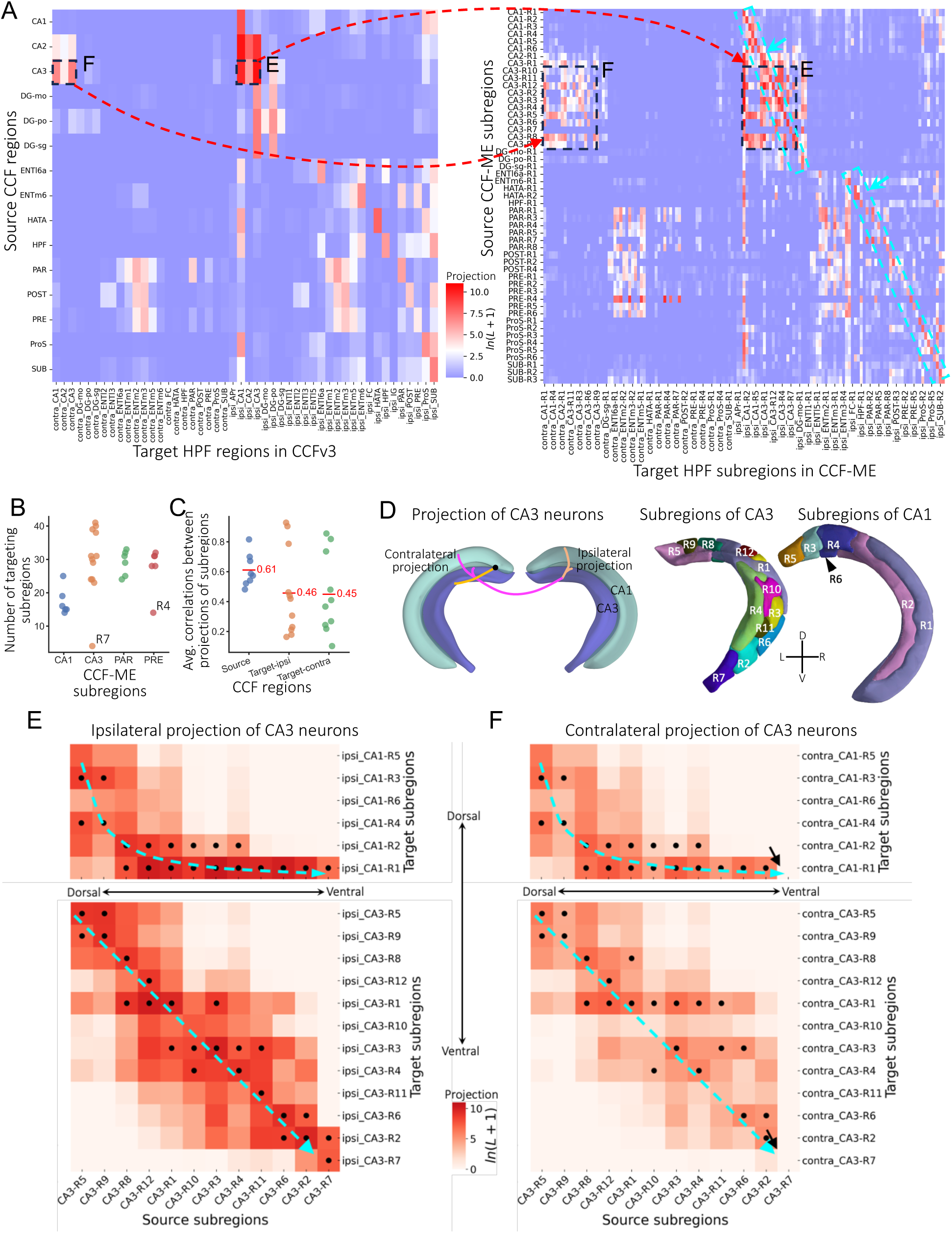
Axonal projection specificity of hippocampal neurons in the CCF-ME Atlas. **A.** Projection matrix of hippocampal neurons across CCF-ME subregions in HPF, HY, and STR. The dashed black rectangles highlight the ipsilateral and contralateral projections of CA3 neurons to CA1 and CA3, which will be discussed in panels D and E. The dashed cyan rectangles illustrate the intra-subregion projections. **B**. Strip plot showing the distribution of the number of targeting subregions for CA1, CA3, PAR, and PRE subregions. Subregion R7 of CA3 and subregion R4 of PRE are explicitly annotated. **C**. Strip plot illustrating the distributions of average correlations between projections of subregions within each CCF region. “Source”, “Target-ipsi”, and “Target-contra” represent the source CCF regions, ipsilateral target regions, and contralateral target regions, respectively. **D**. Schematic illustration of Schaffer collateral projections from CA3 neurons to CA1 (left), the sub-parcellations of CA3 (middle) and CA1 (right). Subregions are distinguished by different colors. D, V, L, and R: dorsal, ventral, left, and right. **E**-**F**. Ipsilateral (E) and contralateral (F) projection matrices of CA3 neurons. Subregions are sorted according to their locations from dorsal to ventral. Dots mark the top two subregions with the highest axonal skeleton lengths in these subregions. Dashed cyan lines highlight the trace of the top subregions from dorsal to ventral. Black arrows indicate the absent contralateral projections from the ventral subregion CA3-R7.

Quantitatively, the number of projected hippocampal formation (HPF) subregions varied across subregions within the same CCF region, as demonstrated by the dispersed numbers in CA1, CA3, PAR, and PRE subregions (**Figure 4B**). A typical CA1 subregion targeted approximately 20 HPF subregions, while subregions in CA3, parasubiculum (PAR), and PRE were projected to roughly 30 subregions, with moderate diversity ranging from 20 to 45 (**Figure 4B**). The R7 subregion of CA3 and the R4 subregion of PRE exhibited notably higher specificity (small numbers of targeted subregions; **Figure 4B**). Correlation analyses further confirmed this specificity, with the average correlation between projections in source subregions and target subregions being 0.61 and ∼0.45, respectively (**Figure 4C**).

The increased specificity in projection mapping enables a more detailed characterization of the Schaffer collateral projections of CA3 neurons to CA1 (Andersen et al., 1971) in both the ipsilateral and contralateral hemispheres, traversing the adjacent CA3 regions. In the CCF-ME atlas, CA3 is subdivided into 12 subregions, primarily along the longitudinal axis. Conversely, CA1 is partitioned into four longitudinally distributed subregions at the dorsal end and two medial-lateral subregions at the ventral end (R1 and R2; **Figure 4D**). The projection matrices reveal that CA3 neurons exhibit nearly parallel projections within CA3 on the ipsilateral side, with the projections predominantly aligned along the diagonal, with limited spatial spreads (**Figure 4E**). Ipsilateral projections from CA3 to CA1 also display this spatial pattern, with neurons in ventral CA3 subregions projecting to ventral CA1 subregions R1 and R2, and dorsal CA3 subregions projecting to dorsal CA1 subregions (**Figure 4E**).

Contralateral projections of CA3 neurons display similar projection patterns but with reduced overall projection strength (**Figure 4F**). A notable observation is that neurons in the extreme ventral subregion R7 of CA3 project exclusively to the ventral subregions of both CA3 and CA1 in the ipsilateral hemisphere, but with few contralateral projections (highlighted by black arrows in **Figure 4F**).

In summary, compared to CCF, the CCF-ME atlas improves the specificity of projection mapping of hippocampal neurons, and illustrates a collateral and spatially correlated projection style from CA3 to both CA3 and CA1.

### Microenvironments capture long-range connectivity specificity of CP neurons without reconstruction

In addition to linking dendritic microenvironments with the axonal projection patterns of hippocampal neurons, we investigated the connectivity specificity of the CCF-ME atlas and compared it with previous studies. First, we validated the sub-parcellation of CP by comparing it to previous parcellations derived from in-CP projections of cortical neurons (Hintiryan et al., 2016; Gao et al., 2022) and diverse projections of CP neurons (Foster et al., 2021). Subsequently, we examined the subregion-specific input and output connectivity patterns of CP using a whole-brain single-neuron connectome.

Similar to the microenvironment features observed in the hippocampus (**Supplementary Figure S5B**), CP neurons also exhibited multidimensional spatial differentiation (**Supplementary Figure S8A**). However, the boundaries between subpopulations were less clear than those in the hippocampus (**Supplementary Figure S5**). Along the anterior-posterior axis, dorsal microenvironments displayed a gradual transition of morphologies to green colors (dorsal sides of slices S1 to S6; **Supplementary Figure S8A**). Subpopulations also emerged along the medial-lateral axis within coronal slices, such as the pink-colored subpopulations in slices S3, S4, and S5 (**Supplementary Figure S8A**). Moreover, morphological differentiation along the dorsal-ventral axis was observed, as dorsal neurons in slices S4, S5, and S6 exhibited varied features compared to ventral neurons (**Supplementary Figure S8A**).

The differentiation of dendritic microenvironments and the resulting sub-parcellation align with the subdomain organization of the CP reported in studies on the cortico-striatal connectome and cortico-basal ganglia-thalamic networks (Hintiryan et al., 2016; Foster et al., 2021). Specifically, subregions in slice S1 are similar to those in the rostral level defined in (Hintiryan et al., 2016), while subregions in slices S3 and S4 align with those in the intermediate level, and subregions in the posterior slices S5 resemble those in the caudal level (**Figure 5B**; Hintiryan et al., 2016). Previously, CP had been categorized into four levels along the anterior-posterior axis: rostral (CP.r), intermediate (CP.i), caudal (CP.c), and extreme (CP.ext; Hintiryan et al., 2016). We extended this classification to five levels by adding a rostral intermediate (CP.ri) level to better describe the cross-slice 3D subregions, and mapped all 13 subregions into these five levels (**Figure 5C**).

**Figure 5.**
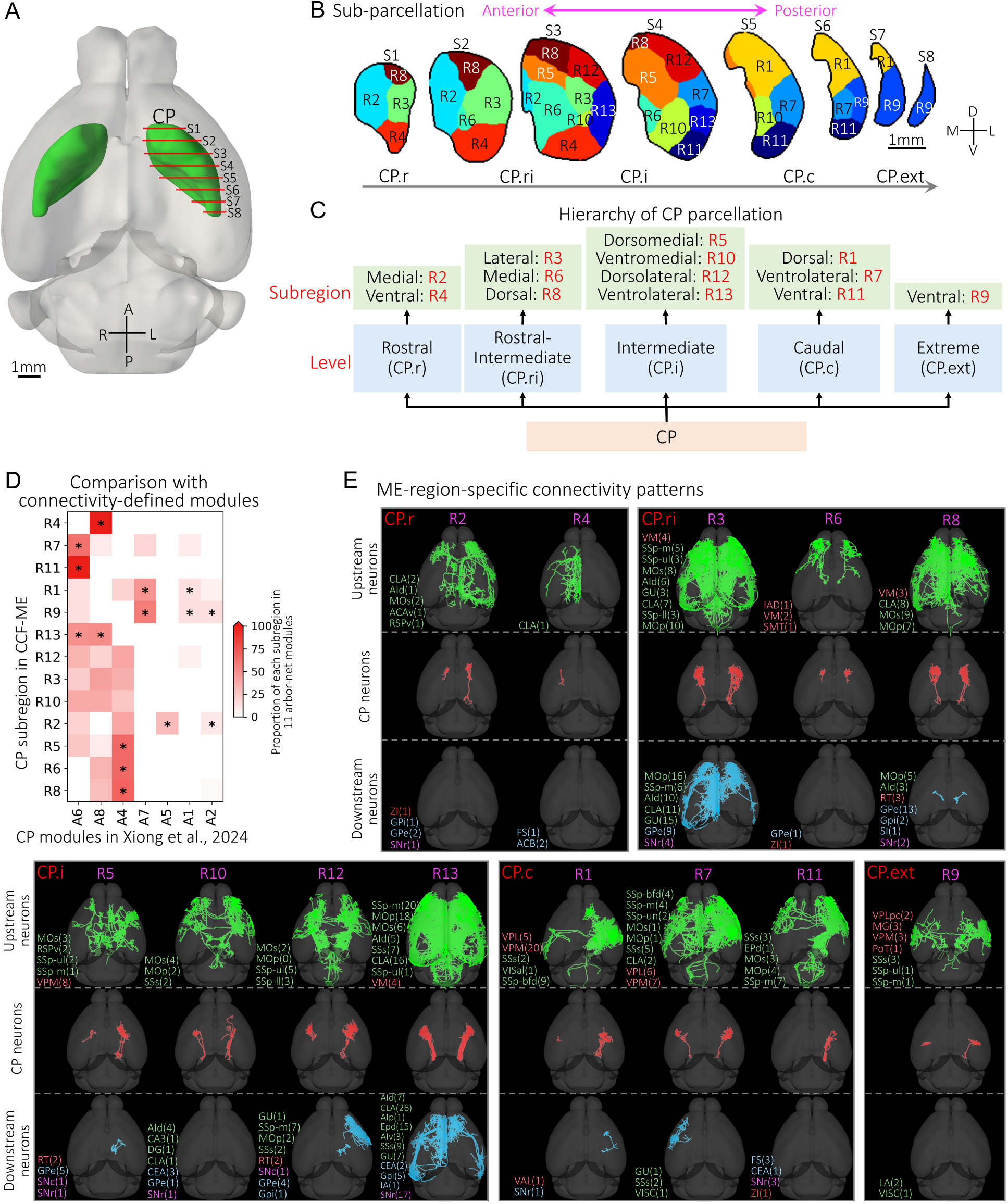
Sub-parcellation of CP and the diversity of neuronal connectivity of CP subregions. **A.** Illustration of CP in the CCFv3 atlas, with eight evenly spaced slices (S1 to S8) from anterior to posterior highlighted by red lines. The distance between two consecutive slices is 0.5 mm. A, P, L, and R: anterior, posterior, left, and right. **B**. Sub-parcellation of CP. The parcellated subregions are shown on the eight slices. Subregions are randomly colored. D, V, M, and L: dorsal, ventral, medial, and lateral. **C**. The anatomical hierarchy of CP subregions, following the nomenclature from (Hintiryan et al., 2016). The putative locations of CP levels (CP.r, CP.ri, CP.i, CP.c, CP.ext) are highlighted at the bottom. **D**. The relationship between CP subregions and connectivity-based modules identified in Xiong et al., 2024, with stars indicating regions that have statistically significant correspondences (*p*-value < 0.0001), determined using a one-sided Fisher’s exact test. **E**. Horizontal views of representative upstream neurons, CP neurons, and downstream neurons for CP subregions. Upstream and downstream neurons were identified based on neuronal connectivity. The identified upstream and downstream CCF brain regions and the numbers of neurons in each region are annotated in parentheses. The regions and corresponding neuron counts are color-coded based on their brain areas: cortical regions are green, midbrain regions cyan, thalamic regions tomato, and hypothalamic regions red. Only full morphologies are shown as examples in the atlas. The full names of the region abbreviations can be found in the **Methods** section.

Furthermore, the CP parcellation in the CCF-ME atlas correlates well with modules derived from the whole-brain single-neuron connectome (Xiong et al., unpublished, under review). Most CP subregions in the CCF-ME correspond to specific modules, except for subregions R3, R10, and R12, which display preferences for modules A8, A6, and A4, respectively (**Figure 5D**). Similarly, the striatal subregions in CCF-ME are consistent with those identified based on the in-CP projection patterns of prefrontal cortical neurons (**Supplementary Figure S8B;** Gao et al., 2022).

Connectivity analyses based on 1,877 manually annotated full neuron morphologies and 18,370 dendritic morphologies across the brain reveal subregion-specific patterns in the CP. For example, subregion R6 receives inputs exclusively from the thalamus (**Figure 5E**). In contrast, subregions R2, R4, R10, R11, and R12 (mostly in the rostral and intermediate CP) primarily receive inputs from cortical neurons (**Figure 5E**), with subregion R4 showing high specificity, receiving input exclusively from claustrum neurons (CLA; **Figure 5E**). Other subregions exhibit mixed cortical and thalamic inputs (**Figure 5E**), with input preferences generally consistent with mesoscale connectome analyses, such as the preferences of input motor and somatosensory neurons in R12 and R13, corresponding to the lateral subregions of CP.i (Hintiryan et al., 2016).

Downstream connectivity also exhibited subregion-specific patterns. For example, only cortical neurons were found downstream of subregions R7 and R9 (**Figure 5E**). In subregion R4, all downstream neurons were located in the striatum (**Figure 5E**). Zona incerta (ZI) neurons were identified exclusively in the downstream networks of subregions R2, R6, and R11, located in the medial CP, where they coexisted with neurons from the cerebral nuclei (e.g., GPe, GPi) or midbrain neurons such as SNr (**Figure 5E**). Thalamic neurons were only detected downstream of subregions R1, R5, R8, and R12, which form the dorsal CP (**Figure 5E**). Some subregions, like R8 and R12, contained neurons from a mix of cortical, midbrain, cerebral nuclei, and thalamic regions (**Figure 5E**).

### CCF-ME identifies complementary subregions of a MERFISH transcriptomic atlas

High-resolution transcriptomic techniques, especially recent spatial transcriptomics like MERFISH (Chen et al., 2015), have revolutionized neuronal cell typing at the whole-brain scale, providing potential fine-grained anatomical characterization. Using MERFISH, comprehensive cell atlases containing over 300 subclasses and 5000 clusters have been released (Yao et al., 2023; Zhang et al., 2023). However, the cell types identified are not always consistent with the existing CCF anatomy. Many region boundaries were missing in the transcriptomic modules, such as the boundary between layer 4 of the primary motor cortex and layer 4 of the primary somatosensory cortex (Zhang et al., 2023).

We compared the CCF-ME with the state-of-the-art MERFISH transcriptomic data (Yao et al., 2023). The clustering of MERFISH transcriptions across CCF-ME regions exhibited a modularized structure (**Figure 6A**). Five modules and eight submodules were identified, encompassing all analyzed regions. Regions within most modules/submodules exhibited high homogeneity concerning anatomical brain areas, as well as the neighborhoods and classes defined in previous work (Yao et al., 2023; **Figure 6A**). Specifically, the majority of cortical regions were clustered in module M1, distinguished by unique neighborhood and class types, except several olfactory regions, such as the main olfactory bulb (**Figure 6A**). Similarly, hindbrain and thalamic regions displayed well-defined modular structures with distinct transcriptomic cell types (M2 and M4; **Figure 6A**). Regions from other brain areas, including the cerebral nuclei and hypothalamus, also exhibited stereotyped patterns (**Figure 6A**). These findings indicate that the CCF-ME is overall consistent with the MERFISH data.

**Figure 6.**
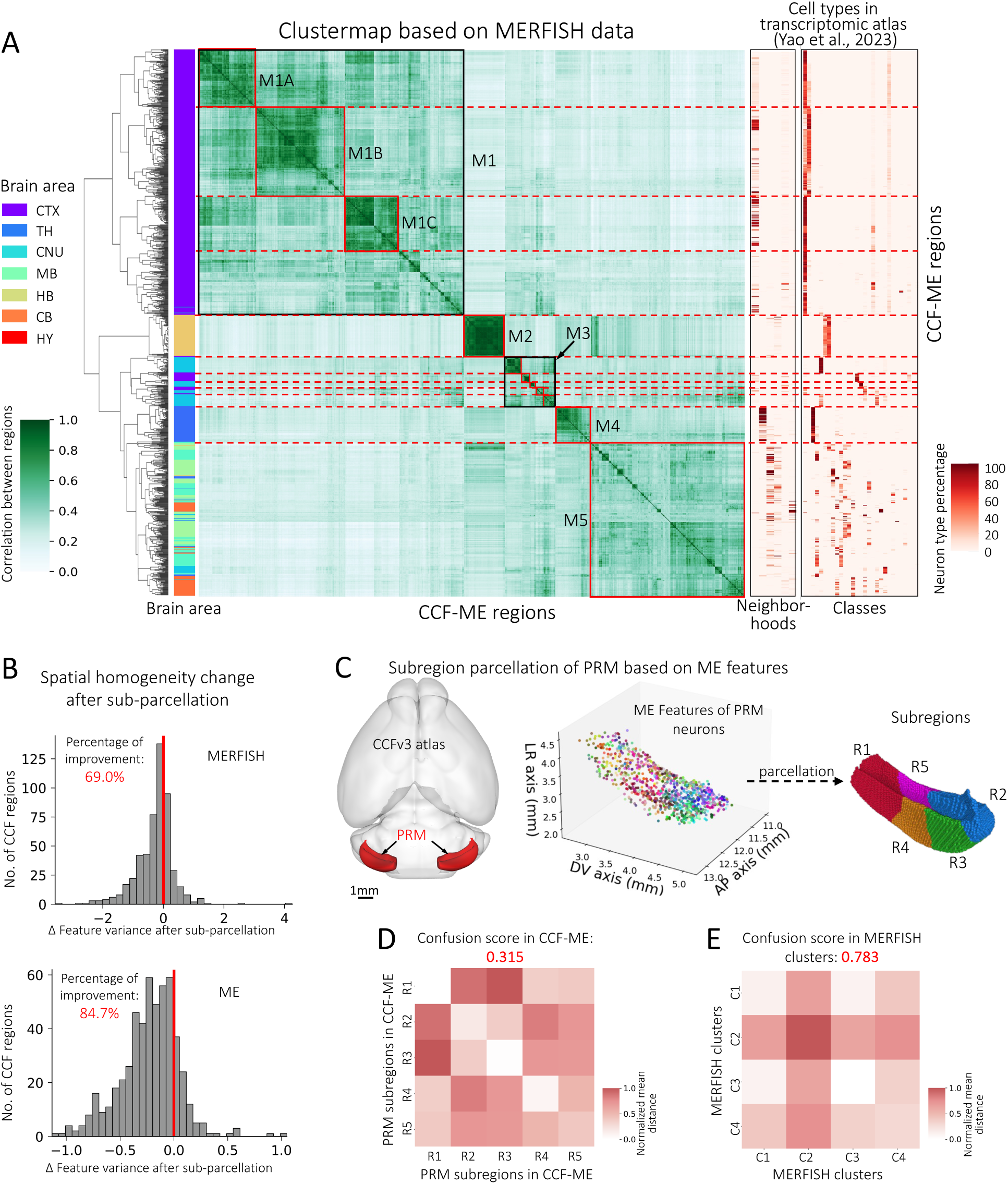
Comparison of microenvironments and MERFISH in identifying subregions. **A.** Clustermap of CCF-ME regions based on MERFISH transcriptomics data. The similarity between two CCF-ME regions is defined as one minus the normalized cosine distance of the average logarithm of transcriptions. Modules (M1, M2, M3, M4, and M5) and submodules (M1A-C, M3A-E) are outlined with solid squares. The dashed red lines highlight the module boundaries. **B.** Histogram showing the variance change in normalized feature space after sub-parcellation for MERFISH data (top) and microenvironments (bottom). The percentages of CCF regions with reduced feature variance (“percentage improvement”) are annotated in red. The red lines indicate the zero-change position, where no change in feature variance occurs. **C.** Sub-parcellation of the CCF region paramedian lobule (PRM) based on microenvironment features (ME features). The ME features are colored according to the top three features. R1, R2, R3, R4, and R5 are the five identified subregions. **D.** Pairwise distance matrix of the parcellated subregions. The distance is the normalized average distance between microenvironment features of all neuron pairs from a pair of regions. The confusion score is highlighted at the top. **E.** Pairwise distance matrix of the four MERFISH clusters of PRM. The distance between two clusters is the normalized average Euclidean distance between all neuron pairs from a pair of clusters. The confusion score is highlighted at the top.

Interestingly, we found that 84.7% of CCF regions showed increased spatial homogeneity in the CCF-ME compared to that in the CCFv3 using the microenvironment data, in contrast to only 69% using MERFISH data (**Figure 6B; Methods**). This suggests that the microenvironment representation can potentially detect subregions that complement those identifiable by other methods, such as MERFISH.

To investigate the biological indication of this finding, we further compared the spatial distributions of MERFISH data and microenvironments using the paramedian lobule (PRM) from the cerebellum as an example (**Figure 6C**). The microenvironments exhibited clear differentiation along the longitudinal axis of the PRM (**Figure 6C**). Neurons in each subregion showed smaller pairwise feature distances, except for neurons in subregion R5, which displayed moderate distances with both subregions R1 and R5 (**Figure 6D**). In contrast, all four MERFISH clusters within the PRM were spatially intermingled and homogeneously distributed (**Supplementary Figure S9**), as intra-cluster neuron pairs exhibited similar spatial distances to inter-cluster distances (**Figure 6E**). Quantitatively, the confusion score between the five subregions of PRM in CCF-ME is 0.315, much smaller than the score between the four clusters of MERFISH data (0.783; **Figure 6E**). In summary, the differentiated spatial preferences of microenvironments provide opportunities for higher-resolution anatomical identification.

## Discussion

The proposed CCF-ME atlas has two main advantages. First, it comprises 1,057 salient regions, 80% more compared to the current 3D standard, CCFv3. CCFv3 is the first high-resolution 3D digital mouse brain atlas that integrates multimodal neuronal information. However, many studies have suggested the potential for further sub-parcellation within CCFv3 brain regions, such as the CP (Hintiryan et al., 2016; Foster et al., 2021; Gao et al., 2022). By leveraging dendritic microenvironments—constructed from neighboring neurons—we were able to parcellate regions into subregions, thus forming the CCF-ME atlas. Second, since CCF-ME is built on top of the CCFv3 framework, it inherits the well-established anatomical ontology and functional annotations.

The key technique behind generating the CCF-ME atlas is the microenvironment representation of dendritic arbors. While single-neuron morphology has long been a critical marker for neuronal cell typing (Zeng & Sanes, 2017), it has not been widely used in atlas construction due to the challenges of obtaining large-scale, full neuronal reconstructions. In contrast, reconstructing dendrites is more feasible and reliable. Despite concerns about the ability of dendritic morphology to classify subtypes of cortical GABAergic interneurons (DeFelipe et al., 2013), our work demonstrates that the microenvironment enhances local homogeneity while preserving subregion differentiation within CCF-defined regions (**Figure 2**). This finding is further supported by the clustering of microenvironments in the hippocampus compared to the indistinguishable single-neuron dendrites (**Figure 3**). Our result may also indicate that the integrated information from neighboring neurons could enhance morphological discriminative power.

Microenvironments also helped uncover hidden associations across different neuronal modalities. We found that the intrinsic ME-related cell subtypes exhibit a unique, strong correlation with their respective distal axonal projection patterns, not seen in other previous analyses. The finding was further confirmed by higher projection mapping specificity and connectivity specificity when using the CCF-ME atlas (**Figure 4**; **Figure 5**). These results suggest that studying dendritic microenvironments may offer valuable insights into functionally relevant axonal projection patterns, especially in large mammalian brains where full axonal reconstructions are not easily achievable. Whether the correlation is a universal principle across the whole brain or specific to certain areas, and to what extent these correlations exist in other areas, are intriguing follow-up questions to be clarified.

Spatial transcriptomic analyses have revolutionized systematic cell atlas research. However, analyses of this transcriptomic data revealed that many of these subclasses spatially overlap, precluding spatial sub-parcellation of the mouse brain (**Figure 6**). In contrast, dendritic microenvironments exhibited either non-overlapping distributions or gradient differentiation for many CCF regions, providing a finer granularity of anatomical organization. This discrepancy may be attributed to the inadequate number of transcripts captured in current techniques or the inactivity of many region-specific genes at the current developmental stage. The spatial resolution achieved through dendritic microenvironment analysis highlights the potential for integrating morphological data with transcriptomic information to create a more comprehensive brain atlas. By combining these advanced methodologies, we can achieve a deeper understanding of brain organization and the functional implications of cellular diversity.

## Acknowledgment

We thank Zuo-Han Zhao for assisting in auto-tracing dendritic morphologies; Shengdian Jiang for assisting in post-processing the reconstructions; and Le Gao for providing the striatal subdomain data published in (Gao et al., 2022). This work is mainly supported by a New Cornerstone grant awarded to H.P. This work was also supported by a STI2030-Major Projects Grant No. 2022ZD0205200/2022ZD0205204, and “the Fundamental Research Funds for the Central Universities” (No. 2242023K5005).

## Author contributions

H.P. conceptualized and managed this study and instructed the detailed development of the methods and experiments. Y.L. led the development of the methods and generated results with the assistance of S.Z., Z.Y., and all coauthors. S.Z. post-processed and evaluated the reconstructions. S.Z. conducted the comparison between spatial transcriptomic data and microenvironment atlas, prepared materials for the sub-parcellation pipeline, and compared the subregion organization of striatum to connectivity-based and projection-based parcellations. Z.Y. mapped the reconstructions onto the common brain space, assisted in analyzing the spatial transcriptomic data and refining the sub-parcellation pipeline. F.X. assessed the changes in spatial variances of axons. Y.L. conducted all other analyses. Y.L. and H.P. wrote the manuscript with the input of all authors.

## Data availability

The reconstructed morphologies and CCF-ME atlas are accessible on Zenodo (https://zenodo.org/records/13761460).

## Code availability

The source codes for neuron tracing, evaluation, microenvironment construction, and analyses are available at https://github.com/SEU-ALLEN-codebase/BrainParcellation. The project was built upon our analytical library for neuron morphology *pylib* (https://github.com/SEU-ALLEN-codebase/pylib). Other dependencies are summarized in the *requirement.txt*, which can be easily installed through the Python package installer (PIP). Morphological feature extraction was performed using the “global_neuron_feature” plugin of Vaa3D (versions Vaa3D-x 1.1.4 and Vaa3D 4.001). Skeleton resampling was done utilizing the “resample_swc” plugin of Vaa3D (version Vaa3D-x 1.1.4). Short segment pruning was conducted using the “pruning_swc_simple” plugin of Vaa3D (version Vaa3D-x 1.1.4). Feature selection was conducted using *pymrmr* (version 0.1.1). The clustering of microenvironments for each region was done using the Leiden algorithm (leidenalg; version 0.10.2). Connected component estimation was carried out with cc3d (version 3.12.4).

## Methods

### Nomenclature and the CCFv3 atlas

Anatomical nomenclatures, abbreviations, and ontology of the brain adhere to the CCFv3 standards. Throughout this paper, we used the 25 µm resolution version of CCFv3 (Wang et al., 2020), which includes 671 brain regions. These regions consist of 8 ventricle regions, 81 fiber tract regions, and 582 salient regions. The salient regions, except one unclassified region (*id*=997), are distributed across seven major brain areas: cortex (CTX, *n*=295), cerebral nuclei (CNU, *n*=27), cerebellum (CB, *n*=21), hindbrain (HB, *n*=77), hypothalamus (HY, *n*=50), midbrain (MB, *n*=60), and thalamus (TH, *n*=51).

The full names of the region abbreviations are: CBN, cerebellar nuclei; CBX, cerebellar cortex; CTXsp, cortical subplate; HPF, hippocampal formation; MY, medulla; OLF, olfactory areas; P, pons; PAL, pallidum; STR, striatum; ACAv, anterior cingulate area, ventral part; ACB, nucleus accumbens; AId, agranular insular area, dorsal part; AIp, agranular insular area, posterior part; AIv, agranular insular area, ventral part; AUD, Auditory areas; CA1, hippocampal area, field CA1; CA2, hippocampal area, field CA2; CA3, hippocampal area, field CA3; CEA, central amygdalar nucleus; CLA, claustrum; CP, caudoputamen; DG, dentate gyrus; DG-mo, dentate gyrus, molecular layer; DG-po, dentate gyrus, polymorph layer; DG-sg, granule cell layers; ENTl, entorhinal area, lateral part; ENTm, entorhinal area, medial part, dorsal zone; EPd, endopiriform nucleus, dorsal part; FS, fundus of striatum; GPe, globus pallidus, external segment; GPi, globus pallidus, internal segment; GU, gustatory areas; HATA, hippocampo-amygdalar transition area; HPF, hippocampal formation; IAD, interanterodorsal nucleus of the thalamus; IA, intercalated amygdalar nucleus; IC, inferior colliculus; ICc, inferior colliculus, central nucleus; LA, lateral amygdalar nucleus; LSc, lateral septal nucleus, caudal part; MG, medial geniculate complex; MOB, main olfactory bulb; MOp, primary motor area; MOs, secondary motor area; PAR, parasubiculum; POST, postsubiculum; PoT, posterior triangular thalamic nucleus; PRE, presubiculum; PRM, paramedian lobule; ProS, prosubiculum; RSPv, retrosplenial area, ventral part; RT, reticular nucleus of the thalamus; SI, substantia innominata; SIM, simple lobular; SMT, submedial nucleus of the thalamus; SNr, substantia nigra, reticular part; SSp-bfd, primary somatosensory area, barrel field; SSp-ll, primary somatosensory area, lower limb; SSp-m, primary somatosensory area, mouse; SSp-ul, primary somatosensory area, upper limb; SSp-un, primary somatosensory area, unassigned; SSs, supplemental somatosensory area; SUB, subiculum; VAL, ventral anterior-lateral complex of the thalamus; VM, ventral medial nucleus of the thalamus; VPL, ventral posterolateral nucleus of the thalamus; VPLpc, ventral posterolateral nucleus of the thalamus, parvicellular part; VPM, ventral posteromedial nucleus of the thalamus; VISal, anterolateral visual area; VISC, visceral area; ZI, zona incerta.

### Local morphology tracing

The sparsely labeled fMOST mouse brains and the 182,497 somas are released in our previous work (Peng et al., 2023). Most brains are accessible from the Brain Image Library (BIL, http://www.brainimagelibrary.org). The utilized brains in this work are listed in **Supplementary Table S1**, and detailed meta-information is documented in (Peng et al., 2023).

The preprocessing and reconstruction protocols were similar to those in (Peng et al., 2023), with the following updates: (1) We utilized 16-bit soma-centered highest resolution image volumes of size 1024×1024×256 voxels (*x*×*y*×*z*; ∼236×236×256 µm^3^), compared to the original 8-bit volumes in the second highest resolution. (2) A robust image enhancement algorithm, NIEND (Zhao et al., 2024), was applied prior to neuron tracing. (3) Only All-Path-Pruning ( APP2; Xiao & Peng, 2013) was used for neuron tracing, instead of both APP2 and neuTube (Feng et al., 2015). (4) We optimized the post-processing of the auto-traced morphologies by adopting more conservative pruning criteria with a 45-degree angle and radius ratio of 2. In this way, we generated 179,568 reconstructions.

To obtain homogeneous skeletons, we registered the morphologies to the CCFv3 atlas using mBrainAligner (Qu et al., 2022; Li et al., 2022). The morphologies were spherically cropped around their soma locations with a radius of 100 µm, ensuring each neuron was represented by its local reconstructions within an isotropic sphere. Skeletons that disconnected to their somas after cropping were removed. To standardize, we resampled the skeletons to ensure that the distance between two successive nodes on the same branch was 2 µm.

We filtered these reconstructions using two criteria: (1) The soma should be located in the 582 salient brain regions. (2) A good reconstruction must have similar morphological features to the feature distributions of manually annotated morphologies (Peng et al., 2023). To do so, morphologies that fell outside the 5% range of the minimum or maximum feature values of “Total Length” and “#Bifurcations” were discarded. The minimal and maximal feature values were calculated separately for different brain areas (e.g., cortex). As such, we finally got 101,136 auto-traced morphologies.

### Morphological feature extraction

Each neuron was represented by a 24-dimensional morphological feature vector. The first 18 features were calculated using the “global_neuron_feature” plugin of Vaa3D (Peng et al., 2014; Liang et al., 2023). These features are: “#Stems”, “#Bifurcations”, “#Branches”, “#Tips”, “Overall Width”, “Overall Height”, “Overall Depth”, “Total Length”, “Volume”, “Max. Euclidean Distance”, “Max. Path Distance”, “Max. Branch Order”, “Avg. Contraction”, “Avg. Fragmentation”, “Avg. Parent-daughter Ratio”, “Avg. Bifurcation Angle Local”, “Avg. Bifurcation Angle Remote”, and “Hausdorff Dimension”. Six other features were: the three values of the first principal component of the PCA analysis of all skeletal nodes (PC11, PC12, PC13), and the variance ratios of the three principal components. Each feature was Z-score standardized separately when calculating distances or constructing a microenvironment.

### Morphology evaluation

Auto-traced local dendritic morphologies were validated in two ways. Firstly, we calculated a commonly used distance metric, bi-directional structure distance, between the traced morphologies and the dendrites of manually annotated morphologies within the same 100 µm spherical range. This metric estimates the average or median distance of all the nearest corresponding points (**Supplementary Figure S1F**).

Secondly, we compared the morphological features of auto-traced morphologies to those of annotated morphologies. To facilitate intuitive comparison, we calculated the relative feature values for all features, where a well-aligned reconstruction would show feature values close to 1. We then plotted a box plot with their statistics and outliers highlighted (**Supplementary Figure S1E**).

Additionally, the quality of the reconstructions was evident in three ways: (1) Morphological features were relatively homogeneous within local neighborhoods but distinct across different regions and areas; (2) The microenvironment features of hippocampal neurons were consistent with those of manually annotated neurons in co-localized regions; (3) Sub-parcellations of striatum derived from the reconstructions aligned well with subdomain organizations reported in various studies.

### Microenvironment construction

A microenvironment is an ensembled representation of a target neuron and, at most, five spatially nearby neurons. For each target neuron, we first extracted all neurons within a local sphere (*R*=166.36 µm) in the CCF space. The radius *R* was the 75th percentile of distances between the sixth nearest neuron and the target neuron for all neurons. If more than six neurons fell within radius R, the six neurons with the smallest feature-space distances to the target neuron were selected. Otherwise, we kept all neurons within the distance *R*.

The construction of a microenvironment was done in the feature space. The feature vectors of neurons from the same microenvironment were integrated through spatial weighting. Specifically, each feature vector was weighted by the exponential of the negative relative distance to the target neuron (**Figure 4B**; Eq. 2).

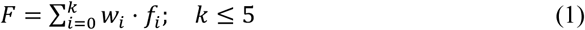

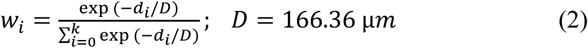

Where *d_i_* is the distance between the *i*-th neuron and the target neuron, *f_i_* is the morphological feature vector, and *k* is the total number of neighboring neurons for the target neuron.

### Subregion parcellation

CCF brain regions were parcellated into subregions based on the spatial organization of microenvironment features. Regions with fewer than 40 neurons or smaller than 225 µm in any direction were skipped. For each eligible region, we constructed an undirected graph based on the spatial distances between microenvironments. Given that many regions are layer-differentiated, we rescaled the spatial distances according to the shape of each region when necessary quantified using principal component analysis (PCA) of all voxels in the region. Edge weights were calculated as the exponential of the negative distance between the feature vectors of connected neurons. All nodes (microenvironments) in the graph were classified using the Leiden algorithm, followed by a majority voting community reassignment among the five nearest microenvironments.

To obtain optimal clustering, we applied an adaptive parameterization strategy for community detection in each region based on silhouette scores. Two parameters were considered: the number of neurons used to initialize the graph and whether to apply shape-dependent scaling for laminar subregions. Results indicated that the best parameter combinations varied across regions. Specifically, shape-dependent spatial scaling was adopted for 73% of regions, while the number of neighbors showed greater variability, with 36% of regions utilizing the nearest neighbor count of 250 neurons. The clustering resulted in a 30% decrease in the average standard deviation of features compared to the original features.

The community for each voxel in the region was predicted using nearest-neighbor interpolation. The resulting parcellation was smoothed by two rounds of 3D median filtering with a ball kernel of 5 voxels (125 µm) in radius. Subregions containing fewer than 512 voxels (0.008 mm³) were discarded and reassigned to nearby communities. During this process, we ensured that each connected component was treated as a subregion unless it was not a connected component in the original CCF atlas, thereby maintaining each subregion as an internally connected mask.

### Feature map visualization

We used the top three features of each neuron to intuitively visualize the spatial distribution of microenvironments. The top three features were identified with the minimum-Redundancy-Max-Relevance (mRMR) algorithm, unless stated otherwise. The three features for the whole-brain feature map are the total length of neuronal skeletons (“Total Length”), average straightness of branches (“Avg. Contraction” or “Avg. Straightness”), and average number of compartments within a branch (“Avg. Fragmentation”, corresponds to branch length).

We mapped these features onto the CCF space according to their soma locations. For better visualization, each feature was first standardized using Z-score normalization, and then histogram equalized to enhance contrast and ensure a uniform distribution of intensities across the feature range of [0, 255], allowing for RGB color coding. Following the left-right convention, microenvironments in the right hemisphere were mirrored to the left hemisphere. This process produced a 3D whole-brain feature map with 101,136 color-coded microenvironments. A similar protocol was applied for generating the feature maps of specific regions.

### Hippocampal formation (HPF) slice stretching

To obtain the longitudinal path for each HPF slice, we extracted the skeletons of the mask slices after performing a morphological closing using a square kernel of size 5 pixels (125 µm). The skeletons were pruned by iteratively removing the shortest branches until only one skeleton remained. This skeleton was then extended on both sides until it reached the boundaries of the mask. This method divided each HPF slice into three distinct components: background, dorsal (closer to the center of the bounding box), and ventral (**Supplementary Figure S6**). The leftmost point of the path termini served as the origin of the new longitudinal-stretched coordinate space. To map all the microenvironments to the longitudinal-stretched space, we identified the nearest point on the longitudinal path as the anchor point for each microenvironment. The path distance from the anchor point to the origin point was used as the x-coordinate, and the distance to the anchor point was the y-coordinate. The dorsal part was assigned negative y-coordinates, and the ventral part positive y-coordinates.

The single-neuron reconstructions from HPF were utilized to compare the morphology distribution of manual reconstructions with our microenvironments. The manual reconstructions were released in (Qiu et al., 2024). A total of 3,822 neurons were retained for this comparison after excluding those without reconstructed dendrites and those located outside the eight regions of interest. For the evaluation of projection specificity among CCF-ME subregions, 10,023 neurons were used after removing neurons with more than one soma.

### Axonal projection mapping

To quantify the axonal projection, we calculated the projection as a vector with a dimensionality of *N*, where *N* corresponds to the number of targeted CCF brain regions or CCF-ME subregions. Each element of the projection vector corresponds to the logarithm of the total axonal length within a region or subregion. To avoid errors during the logarithmic transformation, one was added to the total length before applying the log function.

The projection calculation process began with the resampling of neuronal morphologies (stored in *SWC* format) into two µm-spaced skeletons using Vaa3D to ensure uniform sampling of the skeletons. Subsequently, the axonal components were extracted from the resampled *SWC* file. Given the uniform spacing, the total axonal length in each region or subregion can be estimated as the number of skeletal nodes multiplied by the spacing interval (2 µm). After calculating the axonal length in each region, we compiled these lengths into a projection vector, representing axonal projections across regions or subregions.

### Processing of MERFISH data

We utilized the MERFISH dataset (Zhuang-ABCA-1), publicly available at https://cellxgene.cziscience.com/collections/0cca8620-8dee-45d0-aef5-23f032a5cf09. It comprises 2.6 million spatial transcriptomic cells across 147 coronal sections, featuring a panel of 1,122 genes (Zhang et al., 2023). This dataset was classified into four nested levels: 34 classes, 338 subclasses, 1,201 supertypes, and 5,322 clusters, based on the transcriptomic and spatial atlas (Yao et al., 2023).

Clustering of the raw MERFISH data followed the legacy workflow as follows. Preprocessing steps included removing low-quality cells and genes, normalizing total counts per cell, applying log transformation, selecting highly variable genes, and scaling to unit variance. Following principal component analysis (PCA) and the construction of a neighbor graph, we generated UMAPs and performed Leiden clustering with a resolution parameter set to 0.2.

To assess spatial coherence within brain regions, we utilized two metrics: feature variance and confusion score. Feature variance was calculated by averaging the standard deviations of the top three principal components within a specific brain region, indicating how varied the features were within a specific region. Regarding the confusion score, we first calculated a pairwise distance matrix, where each element represented the average similarity or distance between all intra-subregion or inter-subregion neuron pairs. Then, the confusion score was calculated as the ratio of off-diagonal to diagonal values in the confusion matrix. A lower confusion score reflects greater dissimilarity between subregions. For CCF-ME, the similarity is quantified as the average microenvironment feature distance between neurons. For MERFISH, each element in the matrix represents the average Euclidean distance among neurons within or between clusters (**Supplementary Figure S9**).

### Comparison with connectivity-defined modules of CP

A one-sided Fisher’s exact test was utilized to identify the statistically significant correspondence of CP subregions with the connectivity modules (*p*-value < 0.0001). Major upstream brain regions were identified using a connection score exceeding the 75th percentile based on a dataset comprising 1,877 manually annotated single neuron morphologies (Peng et al., 2023). Downstream brain regions were similarly identified with a connection score greater than 0, utilizing both the 1,877 neurons and 18,370 automatically reconstructed dendritic morphologies.

## Supplementary Figures

**Supplementary Figure S1.**
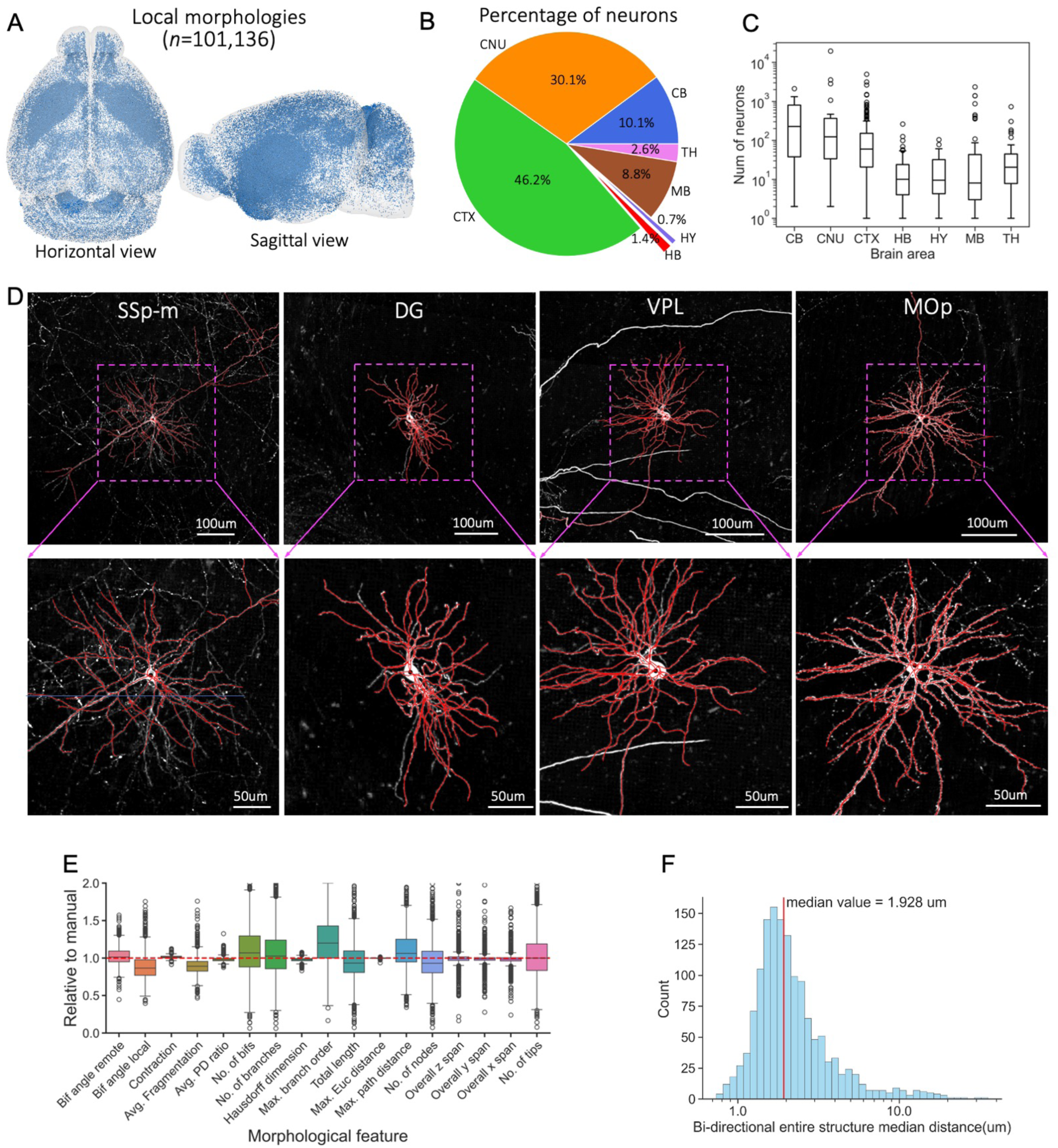
Summary of auto-traced morphologies. **A**. Horizontal and sagittal views of the 101,136 auto-reconstructed neurons mapped in the CCF atlas. **B**. Pie chart showing the distribution of neurons across the brain areas. **C**. Box plot of neuron number distribution within brain regions. **D**. Maximum intensity projections (MIP) of four neuronal images with their corresponding auto-reconstructed morphologies overlaid. The skeletons of each morphology are colored in red. A zoomed-in view of each neuron is shown at the bottom. The soma regions for the neurons are annotated in white at the top. **E**. Box plot showing morphological features of auto-reconstructed neurons relative to those of manually annotated ones. **F**. Histogram of the bi-directional median distance between auto-reconstructed and manually annotated morphologies, with a median value of 1.928 µm.

**Supplementary Figure S2.**
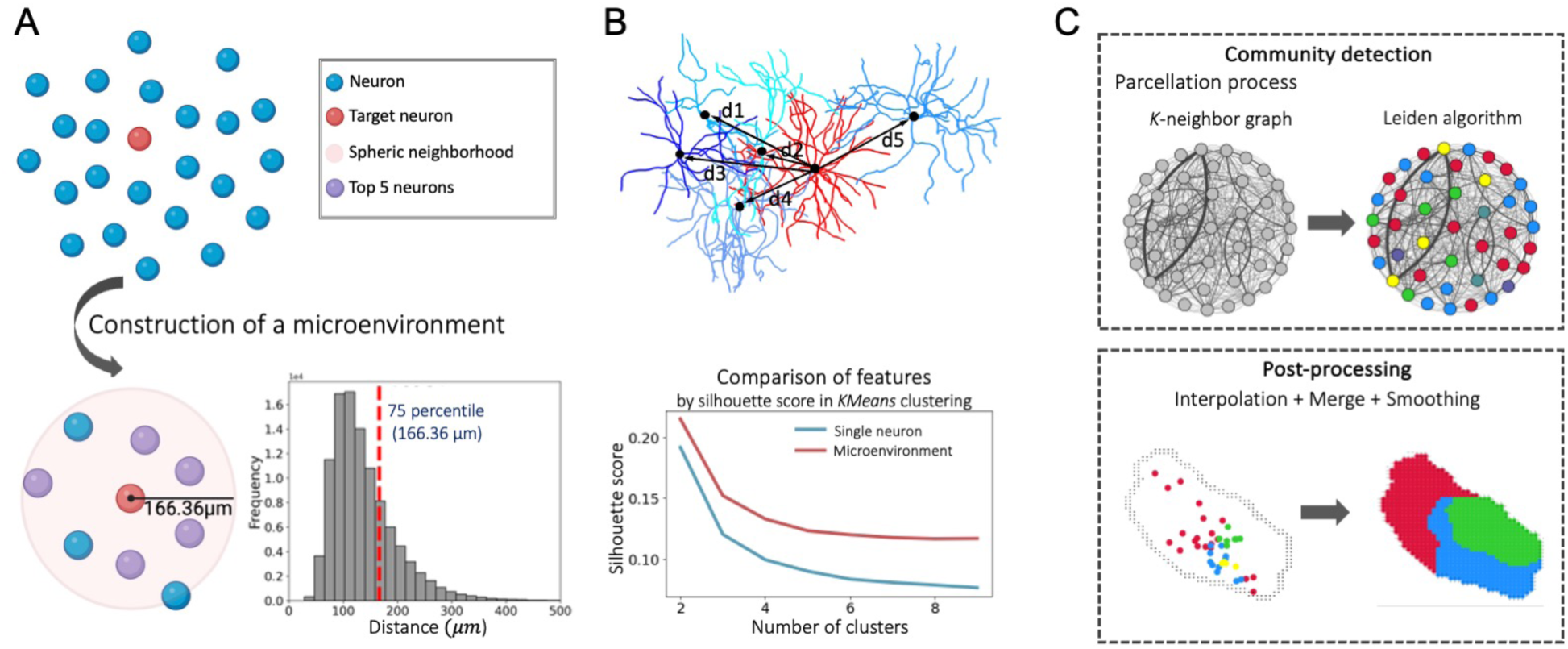
Subregion parcellation based on microenvironments. **A**. Diagram illustrating the construction of a microenvironment. A target neuron (red) is analyzed in the context of its local neighborhood (light pink; bottom). The target neuron identifies its six most similar neighbors (purple) within a 166.36 µm radius (see **Methods**). The histogram shows the frequency distribution of the distances between target neurons and their sixth nearest neurons. **B**. Schematic of the microenvironment feature calculation. A microenvironment is constructed by weighted summarization of the features of the six closest neurons (*i*=0,1,2,3,4,5, where 0 means the target neuron itself). The line plot at the bottom displays the silhouette scores for feature separability of all neurons, as determined by K-Means clustering, across the number of clusters. **C**. Sub-parcellation of brain regions. At the initial community detection step, we construct a K-neighbor graph and use the Leiden algorithm to identify neuron communities. Then, all voxels in the target brain regions are estimated using nearest neighbor interpolation. The resulting subregions are iteratively merged and smoothed to get the final parcellation map. The final parcellation for the example region is illustrated at the bottom.

**Supplementary Figure S3.**
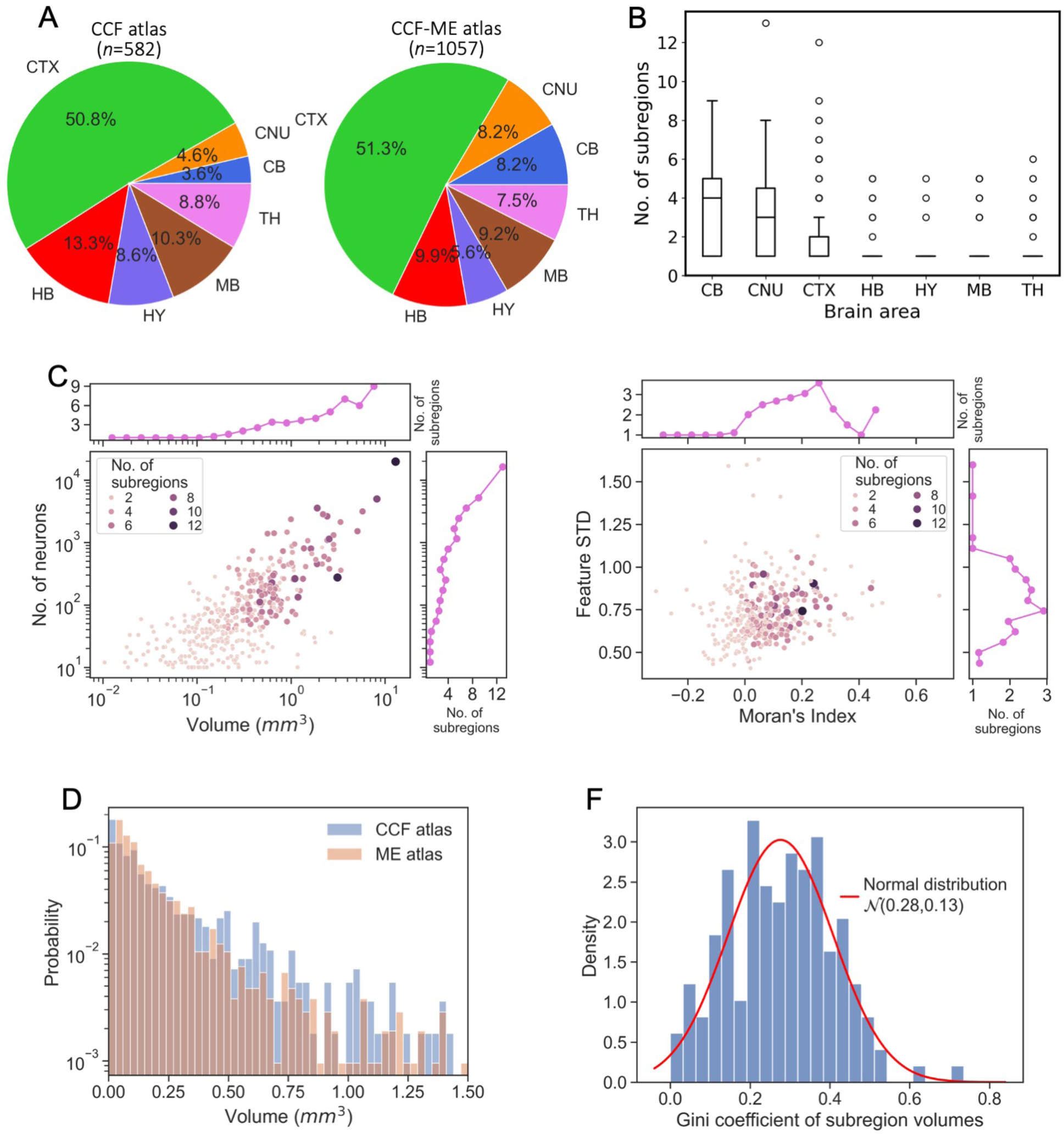
Summary of CCF-ME atlas. **A**. Pie charts showing the percentage of regions in different brain areas in the CCF atlas (*n*=582) and microenvironment atlas (CCF-ME, *n*=1057). **B**. Box plot showing the number of CCF-ME subregions for CCF regions in different brain areas. **C**. Relationship between the number of subregions and four metrics of each CCF region. Left, scatter plot of the region volume (“Volume”) and the number of neurons (“No. of neurons”) against the number of subregions. Both features are plotted in logarithmic space. Right, scatter plot of the standard deviation of microenvironment features (“Feature STD”) and spatial autocorrelation (“Moran’s Index”) against the number of subregions. **D**. Histograms of region volumes in the CCF atlas and ME atlas. The y-axis is presented on an exponential scale for enhanced visualization. **F**. Histogram showing the Gini coefficients of subregion volumes within parcellated CCF regions. A normal density function is fitted (μ=0.28, σ=0.13).

**Supplementary Figure S4.**
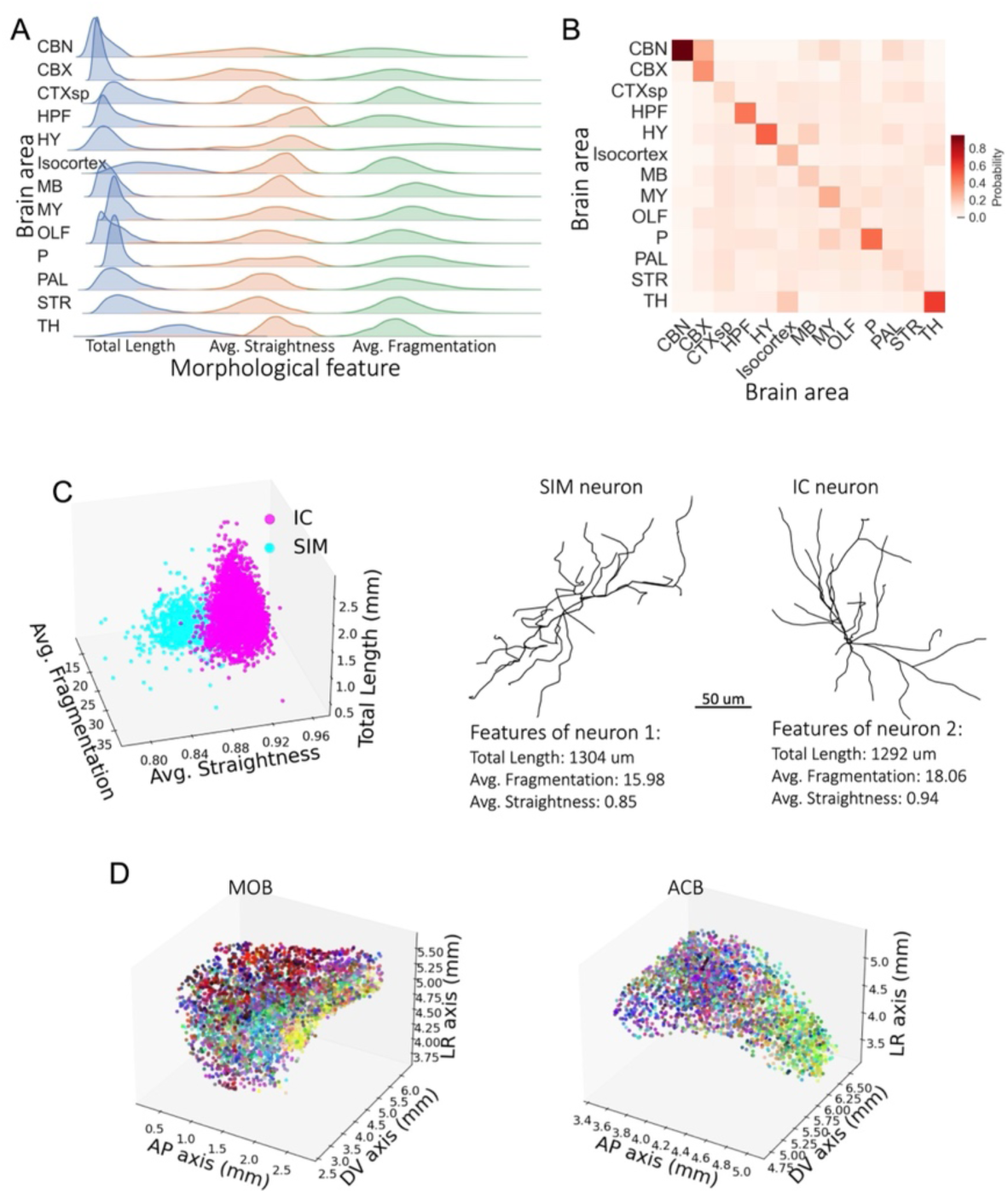
Morphological differentiation across anatomical scales. **A**. Ridge plots showing the distributions of the top three features for 13 brain areas. The abbreviations for these structures follow the CCFv3 nomenclature: cerebellar nuclei (CBN), cerebellar cortex (CBX), cortical subplate (CTXsp), hippocampal formation (HPF), hypothalamus (HY), isocortex, midbrain (MB), medulla (MY), olfactory area (OLF), pons (P), pallidum (PAL), striatum (STR), and thalamus (TH). **B**. Heatmap of the pairwise feature similarity, calculated as the probability that the most similar microenvironment of each microenvironment belongs to a brain area. **C**. Left, the microenvironment feature distribution of the inferior colliculus (IC) neurons and simple lobular (SIM) neurons. Right, skeletal representations of two exemplar neurons from the IC and SIM, respectively, with their top three feature values annotated at the bottom. **D**. Feature landscapes of the main olfactory bulb (MOB) and nucleus accumbens (ACB) in the 3D feature space. The top three features were utilized for visualization. The features were histogram-equalized by channels and were standardized by each region.

**Supplementary Figure S5.**
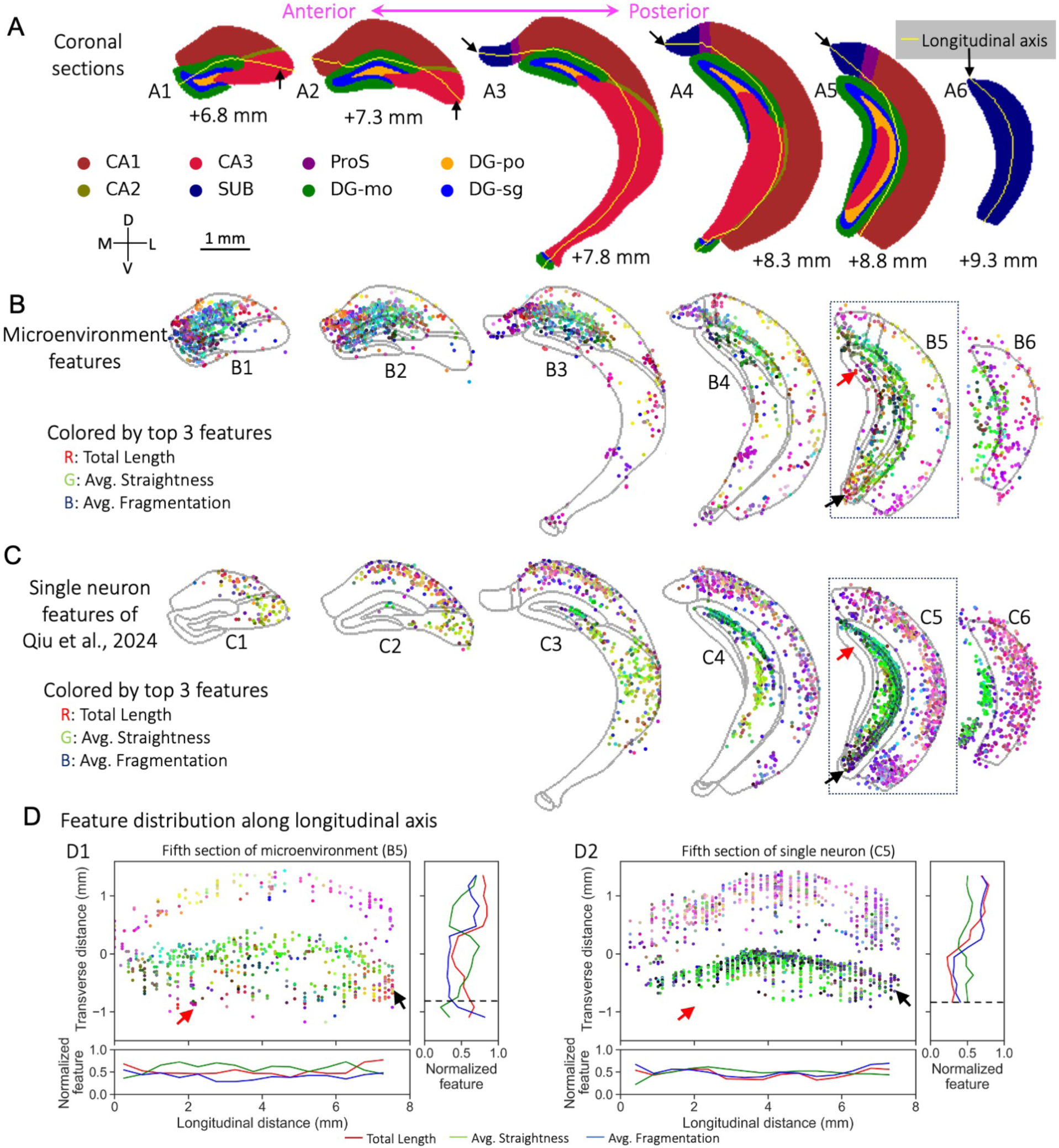
Spatial organization of the hippocampal neurons. **A**. Coronal views of masks for eight hippocampal regions. Six slices ranging from 6.8 to 9.3 mm (from the origin point in the CCF atlas) along the AP axis are shown. The longitudinal axis for each slice is highlighted by a yellow curve and pointed out by black arrows. The transverse axis is perpendicular to the longitudinal axis. D, V, M, and L: dorsal, ventral, medial, and lateral. **B-C**. The top three features of microenvironments (B) and manually annotated single neuron dendrites (C) are projected onto the slices. Neurons within 0.25 mm are mapped to the slice using maximum intensity projection. **D**. Feature distribution in the stretched space for the fifth slice of both microenvironments (left; D1) and single neuron dendrites (right; D2). The bottom and right insets of each component display the mean feature values of microenvironments along the longitudinal and transverse axes, respectively. The horizontal dashed black lines in the right insets indicate the lower limit along the transverse axis for all neurons in the single-neuron dataset.

**Supplementary Figure S6.**
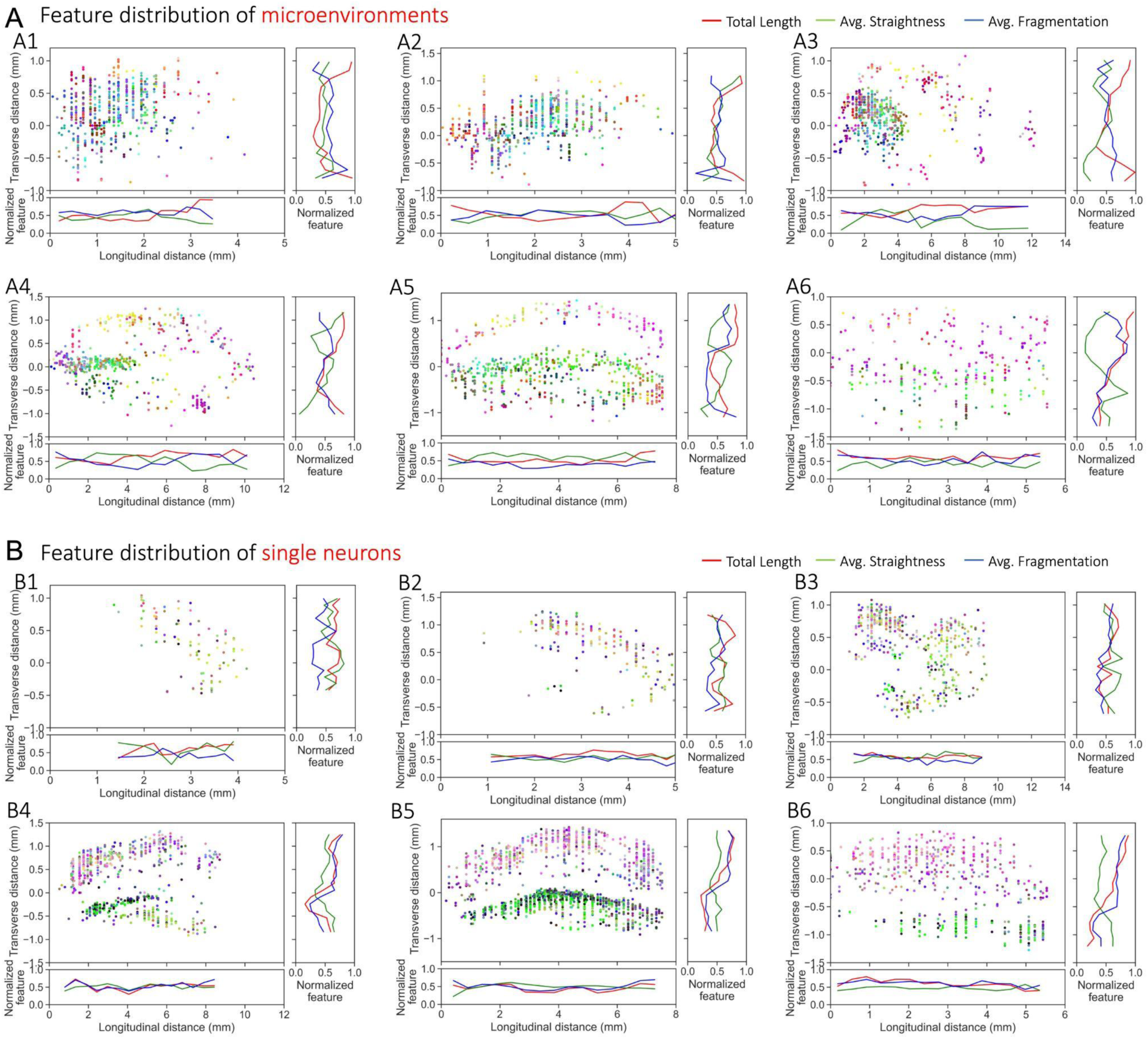
Morphological feature distribution in the stretched space of hippocampal slices. The stretching of a slice is estimated by mapping each microenvironment to a new coordinate system according to the point with the minimal distance along the longitudinal axis (see **Methods**). The bottom and right insets of each component show the mean feature values of microenvironments along the longitudinal and transverse axes. **A-B**. Feature distribution for microenvironments (A) and manually annotated single neuron dendrites (B).

**Supplementary Figure S7.**
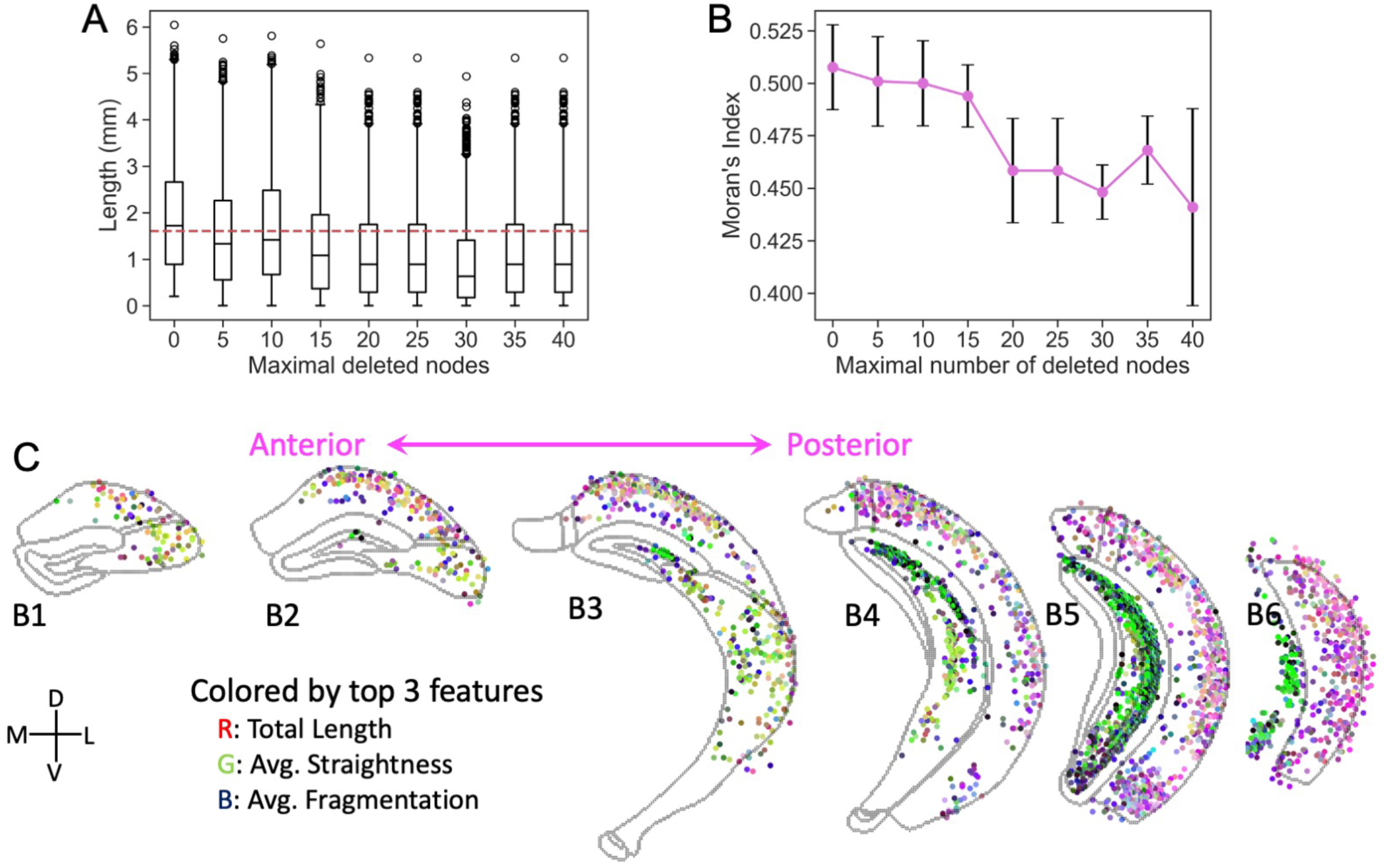
Robustness of spatial organization to morphological perturbations. **A.** Box plots showing the effect of randomly removing skeletal nodes on total skeleton length. The dashed red line indicates the estimated median length of auto-traced local dendrites. **B.** The effect of node removal on spatial homogeneity, measured by Moran’s Index. All downstream nodes connected to the deleted nodes are removed from the skeleton. **C.** Coronal slices showing the spatial distribution of the top three morphological features across the anterior-posterior axis after perturbation by randomly removing up to 40 nodes and all downstream nodes. D, V, M, and L: dorsal, ventral, medial, and lateral.

**Supplementary Figure S8.**
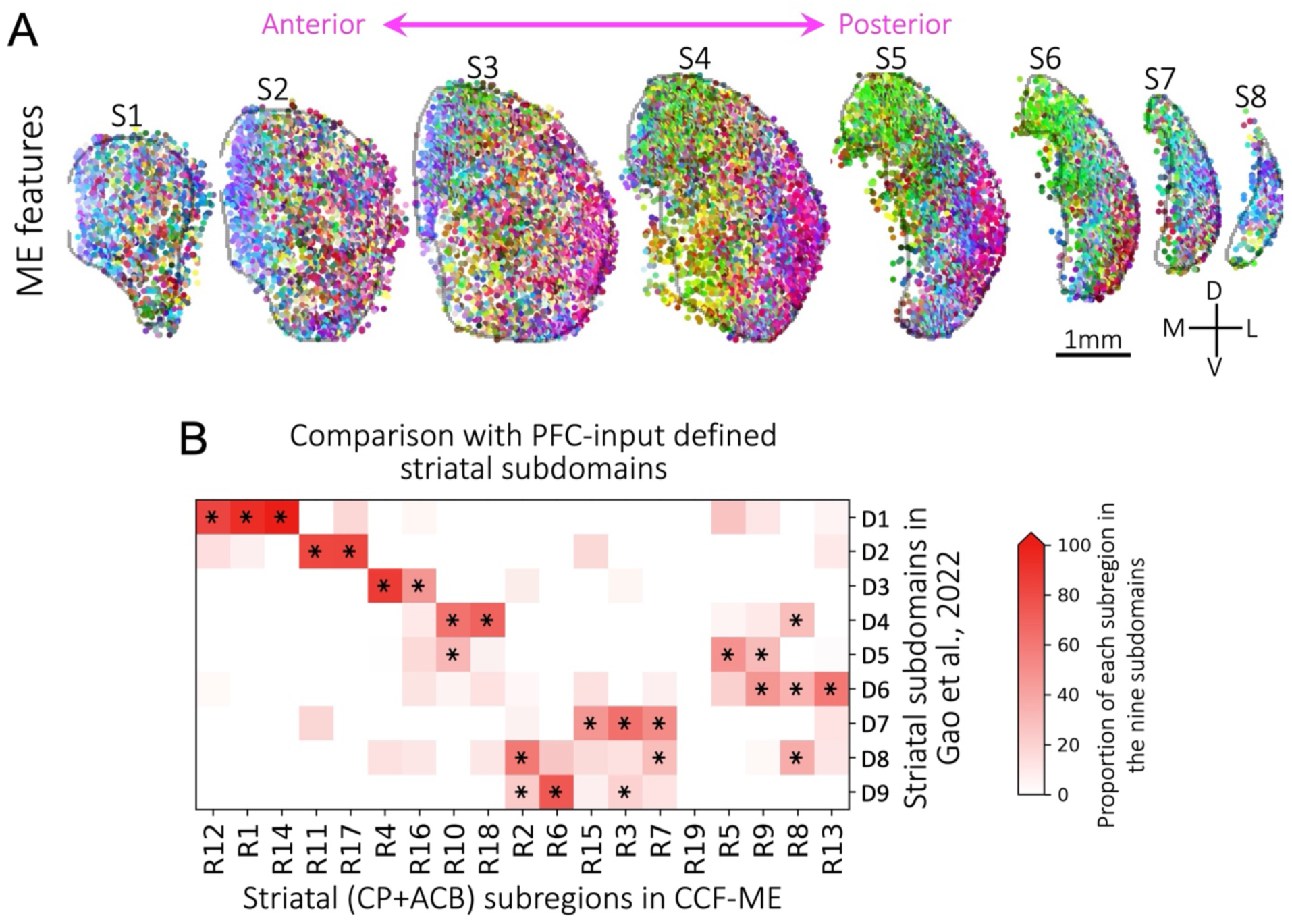
Correspondence between CP subregions in CCF-ME with striatal subdomains or subtypes from other modalities. **A.** Spatial distribution of microenvironmental features for the eight slices. The top three features of the top 24-dimensional morphological features were calculated using PCA and color-coded as red, green, and blue, respectively. D, V, M, and L: dorsal, ventral, medial, and lateral. **B**. The correspondence of striatal subregions with the projection patterns of prefrontal cortical neurons in the striatum is presented. Two CCF regions, CP and ACB, are included for comparison. The stars indicate regions that have statistically significant correspondences (*p*-value < 0.0001), determined using a one-sided Fisher’s exact test.

**Supplementary Figure S9.**
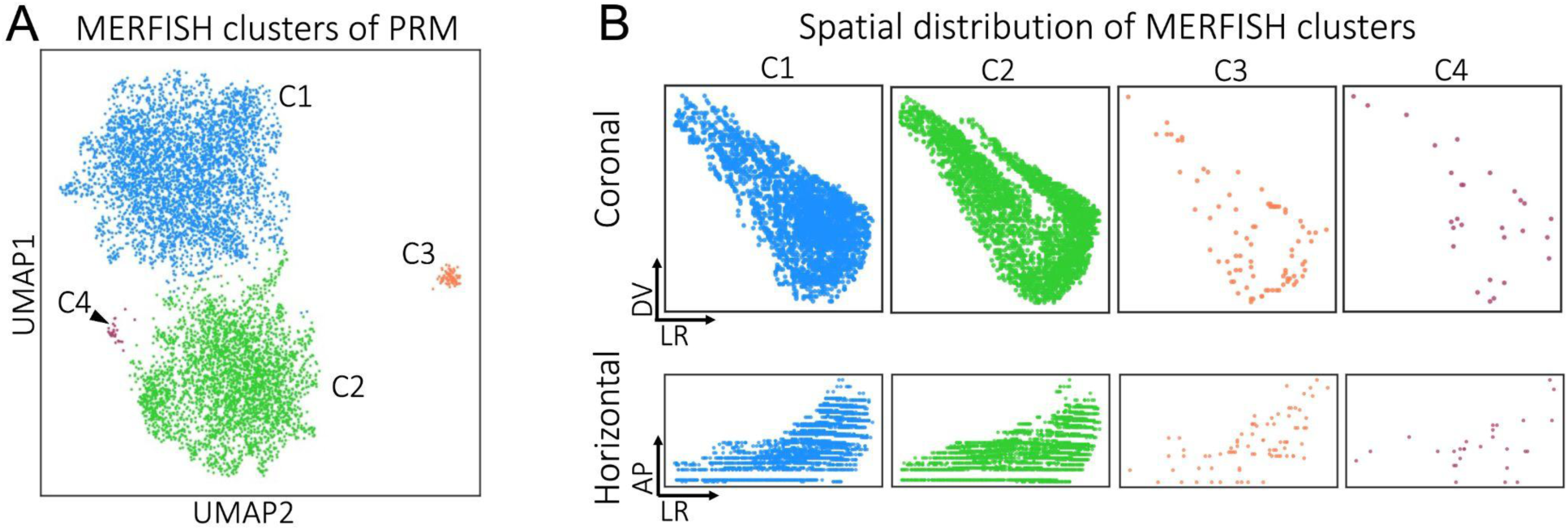
Intermingled spatial distribution of MERFISH clusters of PRM neurons. **A.** Scatter plot showing the four clusters for neurons in PRM based on MERFISH transcriptomic data. **B**. Coronal and horizontal views showing the spatial distribution of the four MERFISH clusters.

## Supplementary Tables

**Supplementary Table S1. The brains utilized in this work.**

